# BLTP2 orchestrates lysobisphosphatidic acid synthesis and exosome biogenesis via SCAMP3-dependent ER contacts in tumorigenesis

**DOI:** 10.1101/2025.04.17.649455

**Authors:** Jingru Wang, Dongchen Li, Yazhou Liu, Tiantian Zhou, Juan Xiong, Wei-Ke Ji

**Affiliations:** Department of Biochemistry and Molecular Biology, School of Basic Medicine, Tongii MedicaCollege and State Key Laboratory for Diagnosis and Treatment of Severe Zoonotic infectiousDiseases, Huazhong University of Science and Technology, Wuhan, China; Department of Ultrasound Medicine, Tangdu Hospital, Fourth Military Medical University, Xi’an, Shaanxi, China; Shenzhen Bay Laboratory, Shenzhen, China; Department of Anesthesiology, Tongji Hospital, Tongji Medical College; Huazhong University of Science and Technology, Wuhan, Hubei, 430030, China; Cell Architecture Research Center, Huazhong University of Science and Technology, Wuhan, China

## Abstract

Multivesicular bodies (MVBs) contain intraluminal vesicles (ILVs) designated for degradation in lysosomes or release as exosomes for cell-to-cell communication. The mechanisms governing ILV/exosome formation are not fully understood. Here, we show that the integral endoplasmic reticulum (ER) membrane protein bridge-like lipid transfer protein 2 (BLTP2; KIAA0100) is indispensable in ILV/exosome formation and that secretory carrier membrane protein 3 (SCAMP3) recruits BLTP2 to ER–MVB membrane contact sites (MCSs) in a Rab5-dependent manner. Our results indicate that this recruitment is hindered by NEDD4-mediated ubiquitination of SCAMP3. Depletion of BLTP2 was found to impede ILV/exosome formation and selectively diminish the levels of cone-shaped phospholipids, including bis(monoacylglycero)phosphate (BMP) and the BMP precursor phosphatidylglycerol (PG) within endosomes. BLTP2 knockout also hampered cell proliferation and tumorigenicity, which could be restored to a significant extent by supplementation with exosomes from wild-type cells. Since BLTP2 is associated with acute monocytic leukemia and is highly expressed in breast cancer, our findings suggest that BLTP2 transfers the BMP/LBPA precursor PG to MVBs for BMP/LBPA synthesis and promotes ILV/exosome formation at SCAMP3-dependent ER–MVB MCSs, a process crucial for cell proliferation and tumorigenesis.

## Introduction

The degradation of harmful and damaged cellular and extracellular material is tightly controlled by endocytic processes, and late endosomes—or multivesicular bodies (MVBs)—play a critical role in the endocytic pathway. They are also vital components of cell-to-cell communication mediated by exosomes. MVBs contain intraluminal vesicles (ILVs) that are formed by the invagination of the endosomal membrane. Cargo is sorted into the ILVs of nascent MVBs, which then mature or detach from endosomes to form free MVBs^1, 2^. These MVBs then deliver the ILVs to lysosomes for degradation or to the plasma membrane, where ILVs are released from the cell in form of exosomes. The degradative and secretory MVB pathways regulate a broad spectrum of functions, and their dysregulation is associated with severe diseases, such as cancer^3–5^.

Endosomal sorting complexes required for transport (ESCRTs) play crucial roles in MVB biogenesis in both the degradative and secretory MVB pathways. In the canonical ESCRT pathway, ubiquitinated cargo and PtdIns3P recruit ESCRT-0 to endosomal membranes for cargo enrichment and recruitment of downstream ESCRTs^1, 6^. ESCRT-I, -II, and -III sequentially mediate the formation and scission of ILVs^1^. Several other factors are also involved in cargo sorting and/or ILV formation, acting in coordination with or independently of ESCRT components^1, 7^. These factors include annexin I^8^, SNX3^9, 10^, SCAMP3^11^, the actin cytoskeleton^12–14^, and certain lipids, such as ceramide^15–17^, phosphatidic acid (PA)^18^, sphingosine-1-phosphate (S1P)^19^, and bis(mono-acylglycero)phosphate^20^, which is also known as lysobisphosphatidic acid (LBPA). Notably, BMP/LBPA is an unconventional phospholipid that is enriched in MVBs/late endosomes and synthesized in late endocytic compartments from phosphatidylglycerol (PG)^21^, an intermediate lipid of the cardiolipin pathway in mitochondria. While recent studies have identified the key enzymes involved in the conversion of PG to lyso-PG and lyso-PG to BMP/LBPA in MVBs/late endosomes (LPLA2^22^ and PLD3/4^23^ or CLN5^24^, respectively), it is not yet clear how PG is transferred from mitochomndria to late endosomes.

In specific cellular contexts^25–29^, including organelle biogenesis^30–32^, organelle trafficking and division^33^, and alleviation of lipotoxicity^34, 35^, the transfer of certain lipid species is mainly mediated by lipid transfer proteins (LTPs) at membrane contact sites (MCSs). LTPs that have a repeating β-groove (RBG) rod-like structure with a hydrophobic channel spanning their entire length have been defined as RGB motif bridge-like lipid transfer proteins (BLTPs)^36, 37^. Recent evidence shows that BLTPs can mediate net lipid transfer to set the stage for the rapid growth of certain organelles^31^. For instance, the BLTP protein Atg2 mediates the transport of phospholipids from the endoplasmic reticulum (ER) to growing autophagosomes to support autophagosome biogenesis during starvation^38, 39^. In addition, Vps13 promotes the formation of the prospore membrane, which encapsulates the daughter nuclei that give rise to spores in yeast^40^. In mammals, Vps13B is required for tubular ERGIC formation^41^ and acrosome biogenesis during sperm development in mice^42^, and Vps13D has been shown to promote peroxisome biogenesis^43^. It has also been shown that SHIP164 regulates endosome–Golgi membrane traffic^44^ and promotes the formation of early endosomal buds^45^. BLTP2 (also known as KIAA0100) is a newly identified BLTP^36, 37^, and has been shown to regulate ciliogenesis^46^ and modulates phosphatidylethanolamine (PE) level in the PM to maintain its fluidity and function^47^. However, its localization in relation to MCS and relavant cellular functions, especially the potential link between BLTP2 and organelle biogenesis, are still largely unknown.

Thus, in this study, we aimed to provide further insight into the mechanisms that support membrane invagination and ILV/exosome formation during MVB biogenesis and to shed light on the role that BLTP2 plays in these processes. We identified KIAA0100/BLTP2 as a novel factor essential for ILV/exosome formation during MVB biogenesis. We further identified SCAMP3 as an adaptor that recruits BLTP2 to ER–MVB MCSs in a Rab5-dependent manner. This recruitment was attenuated by NEDD4-mediated SCAMP3 ubiquitination. Knocking out (KO) BLTP2 blocked ILV/exosome formation and selectively reduced the level of PE, PG, and BMP/LBPA levels in endosomes. BLTP2 KO also impaired cell proliferation and tumorigenicity, which could be restored by the addition of exosomes from wild-type (WT) cells. Hence, we propose that BLTP2 mediates the transfer of the BMP/LBPA precursor PG to MVBs for BMP/LBPA synthesis and ILV/exosome formation at SCAMP3-dependent ER–MVB MCSs and that this process is critical for cell proliferation and tumorigenesis.

## Results

### SCAMP3 is a BLTP2 adaptor that recruits BLTP2 to ER–MVB MCSs

We first investigated the localization of BLTP2 in COS7 cells. We found that BLTP2-GFP was evenly distributed throughout the ER (Fig. 1A). AlphaFold predicted an N-terminal (NT) α-helix in BLTP2, and this could serve as a putative transmembrane (TM) domain (Fig. 1B). A BLTP2 fragment (residues 1– 193) containing the putative TM domain localized to the ER (Fig. 1C), whereas a BLTP2 fragment missing this region did not associate with the ER (Fig. 1D). Therefore, BLTP2 may be an integral membrane protein with an NT TM domain that anchors it to the ER membrane.

**Fig. 1.**
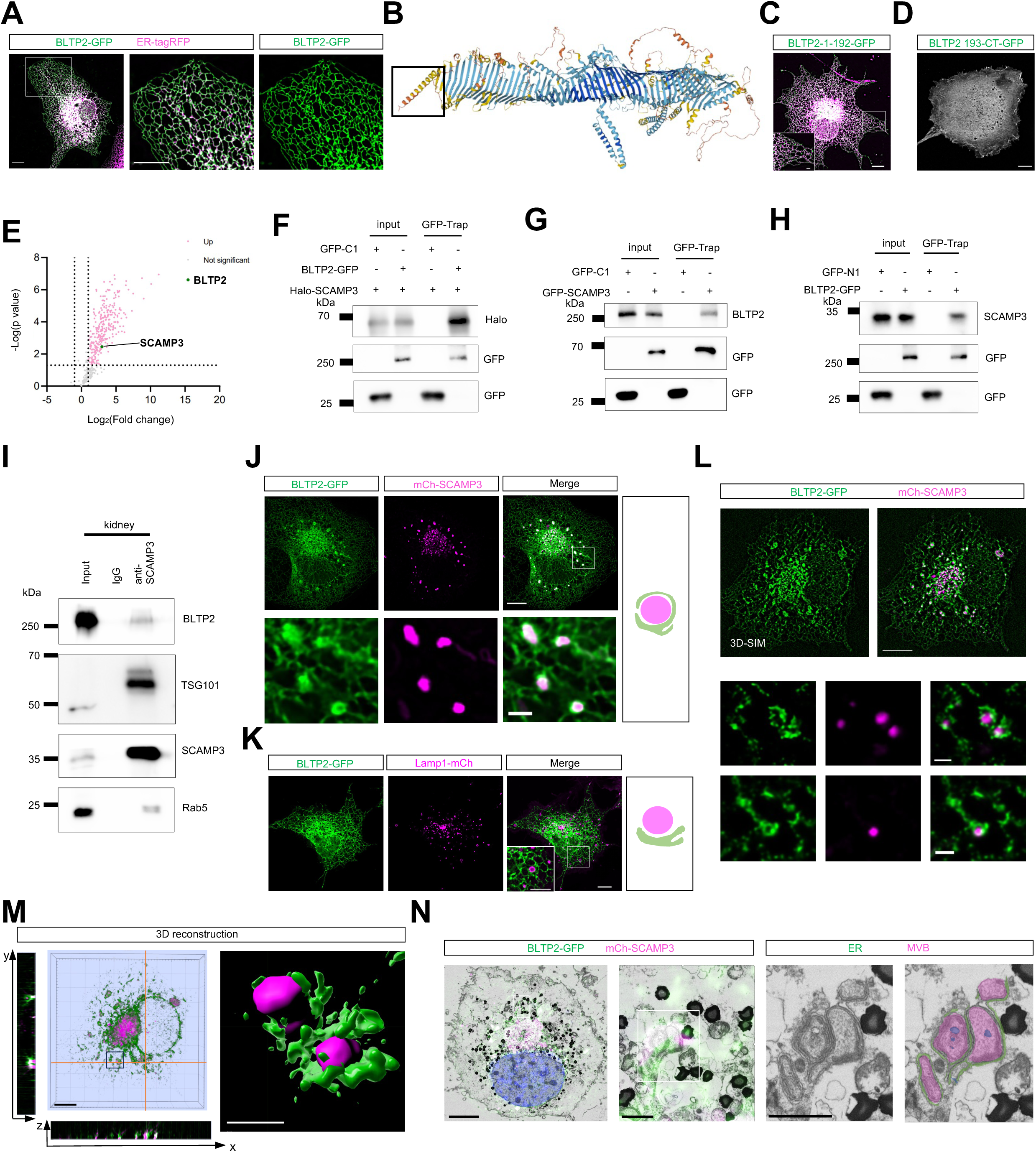
BLTP2 is recruited to ER-MVB MCS by SCAMP3. **A:** Representative images of a live COS7 cell expressing BLTP2-GFP (green) and ER-tagRFP (magenta) with an inset on the right. **B:** The structure of BLTP2 predicted by Alphafold. **C:** Representative images of a live COS7 cell expressing BLTP2-1-193-GFP (green) with an inset. **D:** Representative images of a live COS7 cell expressing BLTP2-193-CT-GFP (grey). **E:** Volcano plot of protein candidates Co-IPed with BLTP2-GFP in HEK293 cells compared with protein candidates coIPed with GFP-tag only. Candidates that were considered significant (-log [P value] > 1.301; P < 0.05) were labeled in pink (Log2 [fold change] > 1; increased in abundance) or gray (Log2 [fold change] <-1; not significant in abundance). **F:** GFP-Trap assays demonstrate interactions between BLTP2-GFP and Halo-SCAMP3 in HEK293 cells. **G:** GFP-Trap assays demonstrate interactions between GFP-SCAMP3 and endogenous BLTP2 in HEK293 cells. **H:** GFP-Trap assays demonstrate interactions between BLTP2-GFP and endogenous SCAMP3 in HEK293 cells. **I:** Co-IP assays show interactions between endogenous SCAMP3 and endogenous BLTP2, TSG101 or Rab5 in mouse kidney lysate. **J, K:** Representative images of COS7 cells expressing BLTP2-GFP (green), along with either mCh-SCAMP3 (**J**, magenta) or Lamp1-mCh (**K**, magenta) or with insets. **L:** Representative 3D-SIM images of a live COS7 cell expressing BLTP2-GFP (green) and mCh-SCAMP3 (magenta) with two insets on the bottom. **M:** Representative 3D rendering of a COS7 cell as in (**L**) with y–z projection to the left and x–z projection to the bottom, with an inset on the right processed by Imaris. **N:** CLEM of a fixed COS7 cell transfected with BLTP2-GFP (green) and mCh-SCAMP3 (magenta) with an inset on the right. Scale bars: 10 μm in the whole cell images and 2 μm in the insets (A, C, D & J-N).

Since BLTP2 is a putative LTP localized to the ER, we hypothesized that it is recruited to an ER-associated MCS via an adaptor resident on the partner organelle. To identify this adaptor, we performed co-immunoprecipitation (co-IP) followed by mass spectrometry (MS) using BLTP2-GFP as the bait. Interestingly, the co-IP–MS analyses showed that the top-ranking BLTP2-interacting proteins were associated with exosomes (Fig. S1A), suggesting a link between BLTP2 and the exosome pathway^48^.

As shown in Fig. 1E, we identified SCAMP3, a tetraspanin integral membrane protein that resides in trans-Golgi vesicles and endosomes and contributes to MVB biogenesis^11^ and sorting events^49^. The results of a series of GFP-trap assays confirmed that both exogenous BLTP2-GFP (Fig. 1F) and endogenous BLTP2 (Fig. 1G) interacted with mCh-SCAMP3 and that endogenous SCAMP3 can be pelleted using BLTP2-GFP (Fig. 1H).

To determine whether BLTP2 interacts with SCAMP3 under physiological conditions, we performed endogenous co-IP assays using SCAMP3 as the bait in mouse kidney tissue lysates. We selected this tissue type because the levels of BLTP2 and SCAMP3 were higher in kidney tissues than in other tissues we tested (Fig. S1B, C). As shown in Fig. 1I, BLTP2 interacted with SCAMP3. In addition, SCAMP3 also interacted with the ESCRT protein TSG101 (consistent with a previous study^49^), and the small GTPase Rab5a.

In contrast to the aforementioned ER localization of BLTP2-GFP (Fig. 1A), mCh-SCAMP3 expression resulted in considerable recruitment of BLTP2-GFP to sites where the ER was juxtaposed with endosomes positive for mCh-SCAMP3, which presumably represented ER–MVB MCSs (Fig. 1J). This recruitment was specific to mCh-SCAMP3; Lamp1-mCh, a general marker for late endosomes or lysosomes, had no effect (Fig. 1K). Super-resolution structured Illumination microscopy (SIM) microscopy was used to confirm the recruitment of BLTP2 by SCAMP3 (Fig. 1L). Notably, the BLTP2 and SCAMP3 molecules appeared to be farther apart at the MCSs than the PDZD8 and Rab7 molecules at the ER–late endosome/lysosome MCSs imaged under the same conditions in our previous study^50^, which is consistent with BLTPs having an extended rod-like structure. In addition, a three-dimensional reconstruction of the SIM images further validated that BLTP2 and SCAMP3 were closely associated, as shown by colocalization in the y–z and x–z projections (Fig. 1M). We also performed correlative light and electron microscopy (CLEM) to analyze the recruitment of BLTP2 by SCAMP3 to ER–MVB MCSs. The obtained images showed that the ER was closely adjacent to mCh-SCAMP3-positive structures that contained several ILVs and presumably represented MVBs (Fig. 1N). Collectively, these results suggest that SCAMP3 recruits BLTP2 to ER–MVB MCSs.

In mammals, the SCAMP family contains four ubiquitous isoforms (SCAMP1–4) and a neuronal isoform (SCAMP5)^51, 52^. Therefore, we investigated whether the other SCAMPs can recruit BLTP2. Flag-tagged SCAMP1, SCAMP2, and SCAMP4 were able to recruit BLTP2; however, SCAMP5 could not recruit BLTP2 (Fig. S1D). In addition, the results of a series of co-IP assays showed that SCAMP1, SCAMP2, SCAMP3, and SCAMP4, but not SCAMP5, interacted with BLTP2-GFP, with SCAMP3 recording the strongest interaction (Fig. S1E).

### An C-terminal (CT) α-helix in BLTP2 is responsible for the interaction with SCAMP3

We then examined how BLTP2 interacts with SCAMP3. Live-cell microscopy revealed that a C-terminal (CT) fragment (residues 1703–1901) of BLTP2 was soluble in the cytosol (Fig. 2A) and strongly recruited by mCh-SCAMP3 (Fig. 2B). In contrast, BLTP2 missing these residues was not recruited by mCh-SCAMP3 (Fig. 2C, D). These results suggest that this region is critical for BLTP2 recruitment by SCAMP3.

**Fig. 2.**
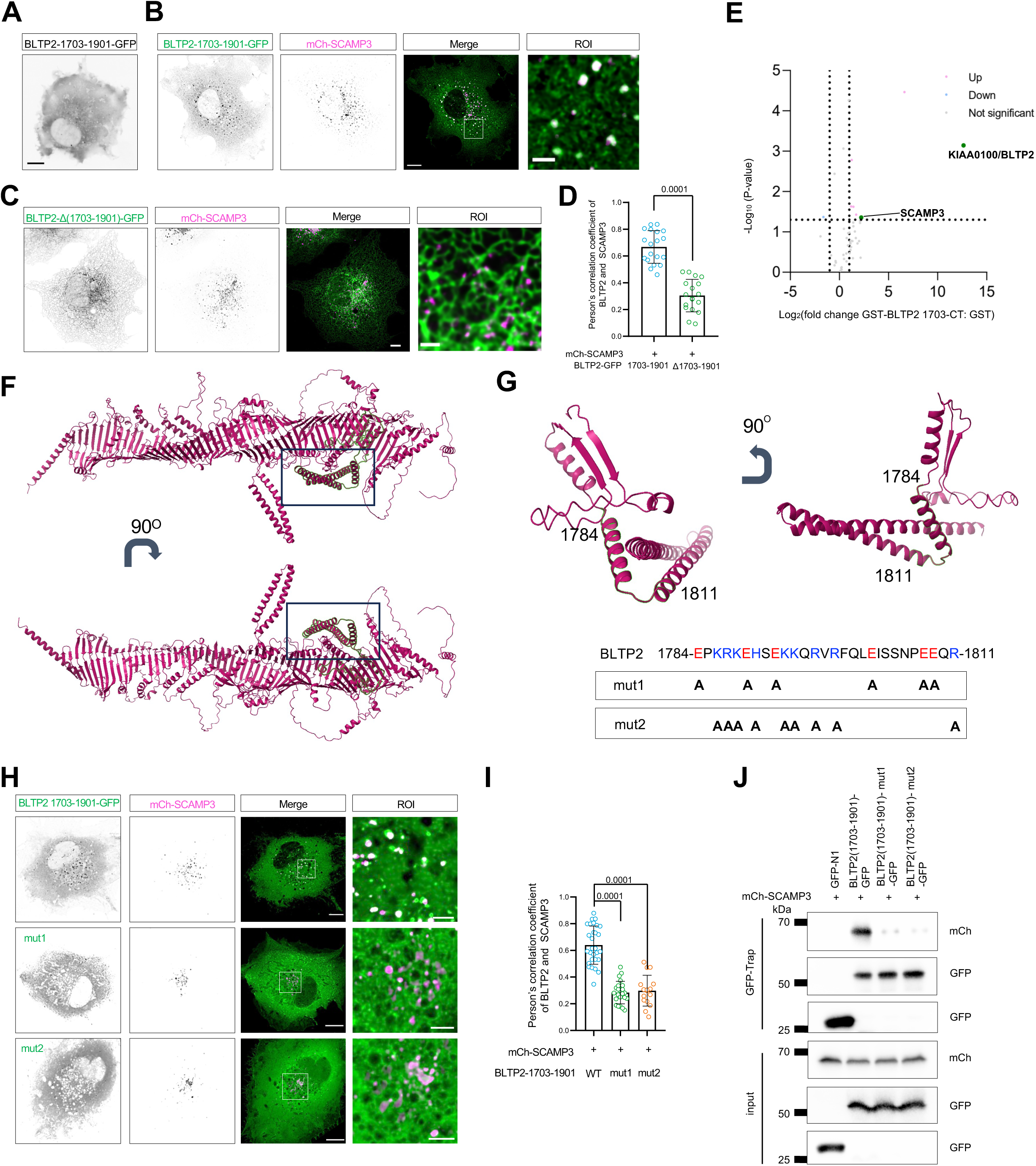
An α-helix in the CT of BLTP2 is responsible for the interaction with SCAMP3. **A**: Representative images of a live COS7 cell expressing BLTP2(1703-1901)-GFP. **B, C:** Representative images of COS7 cells expressing either BLTP2(1703-1901)-GFP (**B**, green) or BLTP2-Δ(1703-1901)-GFP (**C**, green) and mCh-SCAMP3 with an inset on the right. **D:** Pearson’s correlation coefficient of SCAMP3 vs BLTP2(1703-1901) or BLTP2-Δ (1703-1901) in three independent experiments. Two-tailed unpaired Student’s t test. Mean ± SD. **E:** Volcano plot of protein candidates pulldown with BLTP2-1703-1901-GST in mouse kidney lysate. Candidates that were considered significant (-log [P value] > 1.301; P < 0.05) were labeled in pink (Log2 [fold change] > 1; increased in abundance) or blue (Log2 [fold change] <-1; decrease in abundance). **F:** The region containing residues 1703-1901 highlighted in Alphafold predicted BLTP2 structure. **G:** The structure of BLTP2(1703-1901) predicted by Alphafold with two mutants on the bottom. **H:** Representative images of COS7 cells expressing either BLTP2(1703-1901)-GFP (top, green), BLTP2(1703-1901)-mut1-GFP (middle, green) or BLTP2(1703-1901)-mut2-GFP (bottom, green) and mCh-SCAMP3 (magenta) with insets on the right. **I:** Pearson’s correlation coefficient of SCAMP3 vs either BLTP2 (1703-1901), BLTP2(1703-1901)-mut1 or BLTP2(1703-1901)-mut2. More than three independent experiments. Ordinary one-way ANOVA with Tukey’s multiple comparisons test. Mean ± SD. **J:** GFP-Trap assays demonstrate interactions between BLTP2(1703-1901) or its two mutants and mCh-SCAMP3 in HEK293 cells. Scale bars: 10 μm in the whole cell images and 2 μm in the insets (A, B, C & H).

To confirm this notion, we performed pulldown assays with a glutathione S-transferase (GST)-tagged BLTP2 fragment (residues 1703–1901) and mouse kidney lysate, followed by MS. After removing the proteins bound to the GST tag, endogenous SCAMP3 was found to be associated with the BLTP2 fragment and was the second-most abundant protein, as measured by the fold-change ratio (Fig. 2E).

The region of BLTP2 covered by residues 1703–1901 consists of four α-helices and two β-sheets, and the latter elements form part of the extended hydrophobic groove (Fig. 2F). Since three of the four α-helices face outward, we investigated whether any of these helices are required for interaction with SCAMP3, and if so, which helices. Multiple charged residues were found to be present in a small region containing an α-helix and a part of a downstream linker (Fig. 2G, upper panel). Therefore, we hypothesized that these residues may mediate recruitment via electrostatic interactions. To test this hypothesis, we generated two BLTP2 mutants in which these residues were changed to Ala (Fig. 2G, bottom panel) and examined their recruitment by SCAMP3. The imaging results showed that neither mutant was recruited by SCAMP3 (Fig. 2H, G), and the results of GFP-trap assays confirmed that the interactions between SCAMP3 and the mutants were greatly reduced (Fig. 2J). Collectively, these results suggest that the charged residues within the α-helix and the linker are important for the interaction with SCAMP3.

### An α-Helix in the NT of SCAMP3 interacts with BLTP2

Next, we turned our attention to SCAMP3 to determine how this protein recruits BLTP2. SCAMP3 has a large NT domain that contains Asn-Pro-Phe (NPF) repeats and a Pro-rich region, as well as four consecutive TM domains (a tetraspanin region), and a cytoplasmic CT domain^49^ (Fig. 3A). We prepared a series of truncated versions of SCAMP3 and tested their ability to recruit BLTP2 using live-cell microscopy. SCAMP3 without the NPF region (SCAMP3-ΔNPF; Fig. 3B), the Pro-rich region (SCAMP3-ΔP-rich; Fig. 3C), or the CT domain (SCAMP3-ΔCT; Fig. 3D) could still recruit BLTP2. Similarly, the SCAMP3 that was missing a section of the linker between the Pro-rich region and the tetraspanin region could still recruit BLTP2 (SCAMP3-Δ130-156; Fig. 3E). However, deleting another section of the linker (SCAMP3-Δ80-120; Fig. 3F, G) abolished the ability to recruit BLTP2.

**Fig. 3.**
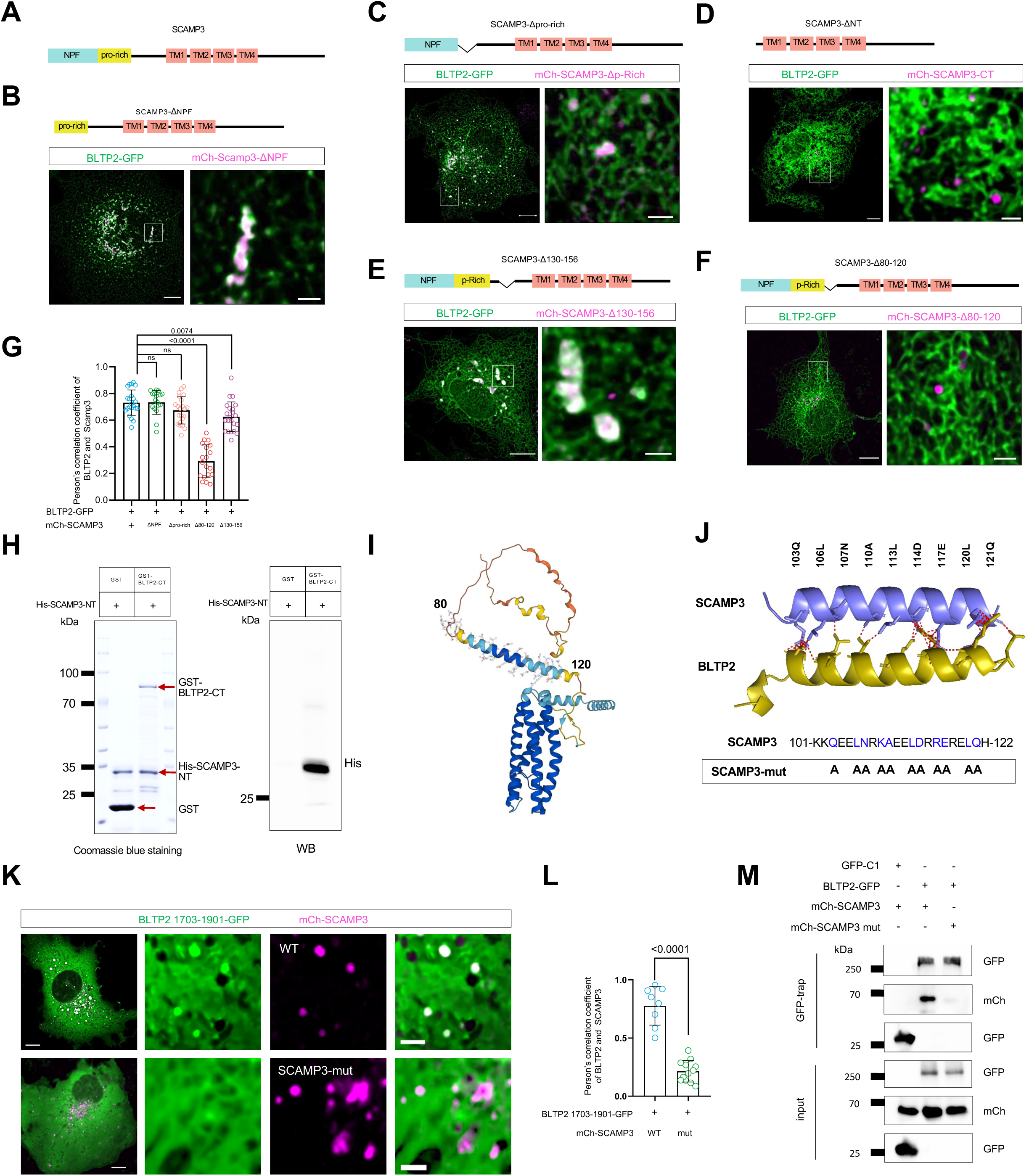
A disorder region in the NT region of SCAMP3 is required for the recruitment of BLTP2. **A**: Domain architecture of human SCAMP3. **B-F:** Representative images of COS7 cells expressing BLTP2-GFP (green) and SCAMP3-ΔNPF (**B**, magenta), SCAMP3-ΔP-rich (**C**, magenta), SCAMP3-ΔNT (**D**, magenta), SCAMP3-Δ80-120 (**E**, magenta) or SCAMP3-Δ130-150 (**F**, magenta) with insets on the right. **G:** Pearson’s correlation coefficient of BLTP2 vs either SCAMP3-ΔNPF, SCAMP3-ΔP-rich, SCAMP3-ΔNT, SCAMP3-Δ80-120 or SCAMP3-Δ130-150. More than three independent experiments. Ordinary one-way ANOVA with Tukey’s multiple comparisons test. Mean ± SD. **H:** GST pull-down assays demonstrate that purified BLTP2-CT (residues 1703-2235), but not GST tag, was pelleted with purified His-SCAMP3-NT (residues 1–170) in vitro. Coomassie blue staining of proteins used in the assay was shown on the left. Red arrows denote the purified proteins in the gel. **I:** The structure of SCAMP3 predicted by Alphafold. **J:** The putative electrostatic interactions between two α-helice of BLTP2 and SCAMP3 with a SCAMP3-mut on the bottom. **K:** Representative images of COS7 cells expressing BLTP2(1703-1901)-GFP (green) and mCh-SCAMP3 (top, magenta), or mCh-SCAMP3-mut (bottom, magenta) with an inset on the right. **L:** Pearson’s correlation coefficient of BLTP2(1703-1901) vs either SCAMP3 or SCAMP3-mut. More than three independent experiments. Two-tailed unpaired Student’s t test. Mean ± SD. **M:** GFP-Trap assays demonstrate interactions between BLTP2(1703-1901) and mCh-SCAMP3 or its mutant in HEK293 cells. Scale bars: 10 μm in the whole cell images and 2 μm in the insets (B-F & K).

Based on these findings, we examined whether SCAMP3 directly interacts with BLTP2 via this region (residues 80–120) by performing in vitro pulldown assays. We used the GST-tagged BLTP2 fragment (residues 1703–1901) to pellet a His-tagged SCAMP3 fragment (residues 80–120; His-SCAMP3-NT). The BLTP2 fragment, but not the GST tag, pelleted His-SCAMP3-NT (Fig. 3H), indicating that the CT of BLTP2 directly interacts with the NT of SCAMP3.

Given that the NT of SCAMP3 consists of an α-helix enriched with charged residues that face the cytosol (Fig. 3I) and that charged residues in the CT domain of BLTP2 were found to be required for the BLTP2–SCAMP3 interaction, we investigated whether the two proteins interact via electrostatic interactions (Fig. 3J, top panel). To test this idea, we changed these charged residues in SCAMP3 to Ala (Fig. 3J, bottom panel) and assessed the ability of the resultant SCAMP3 mutant to recruit BLTP2. The imaging results showed that the SCAMP3 mutant was unable to recruit the CT fragment of BLTP2 (Fig. 3K, L). Consistently, the results of GFP-trap assays showed that the interaction between the CT fragment of BLTP2 and the SCAMP3 mutant was greatly reduced (Fig. 3M). Together, these results suggest that electrostatic interactions between two α-helices mediate the BLTP2–SCAMP3 interaction.

### The recruitment of BLTP2 by SCAMP3 is dependent on Rab5

Next, we investigated the colocalization of BLTP2 and SCAMP3 under endogenous conditions using immunofluorescence (IF) staining. We found that endogenous, untagged BLTP2 formed puncta on the ER (Fig. S2A), which suggested that the even distribution of BLTP2 on the ER resulted from overexpression. The specificity of the BLTP2 and SCAMP3 antibodies used in the IF assay was validated in BLTP2-knockout (KO) HeLa cells prepared using the CRISPR-Cas9 system (clone 5; Figs. S2A, bottom panel, and S2C–G) and HeLa cells in which SCAMP3 was depleted using small interfering RNAs (siRNAs) (Fig. S2H, I).

Importantly, we found that a significant portion (∼45%) of the endogenous BLTP2 puncta were associated with endogenous SCAMP3 (Fig. 4A). However, a fraction (∼42%) of the BLTP2 puncta did not colocalize with SCAMP3 (Fig. 4B), suggesting that the BLTP2–SCAMP3 association may be subjected to regulation.

**Fig. 4.**
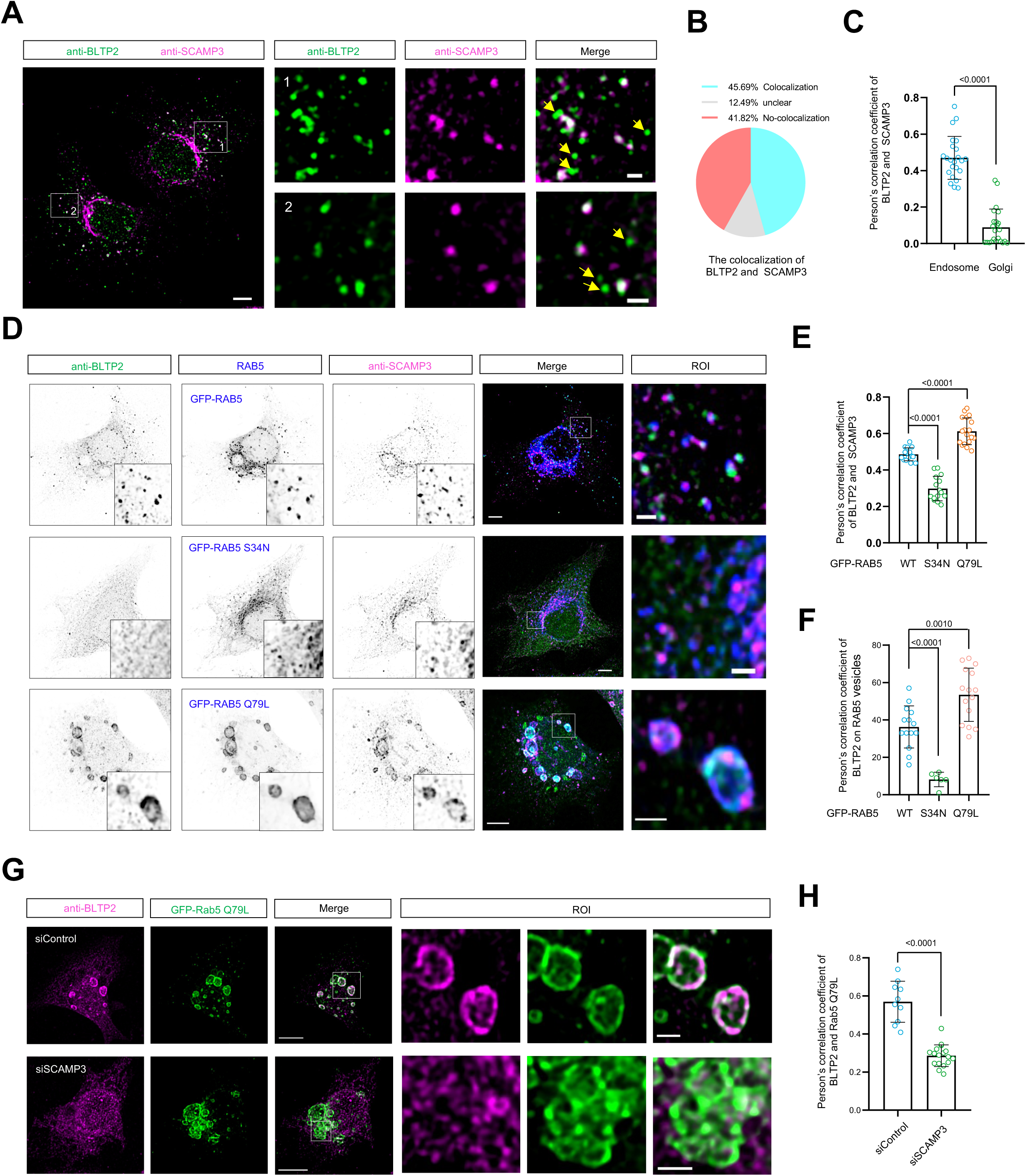
The recruitment of BLTP2 by SCAMP3 is Rab5 activity-dependent. **A:** Representative images of fixed HeLa cells stained with antibodies against BLTP2 (green) and SCAMP3 (magenta) with insets on the right. Yellow arrows denote BLTP2 puncta not colocalizing with SCAMP3. **B:** The colocalization between endogenous BLTP2 and SCAMP3 from more than three independent experiments as in (**A**). **C:** Pearson’s correlation coefficient of endogenous BLTP2 vs endogenous SCAMP3 at endosomal or the Golgi area from more than three independent experiments as in (**A**). Two-tailed unpaired Student’s t test. Mean ± SD. **D:** Representative images of fixed HeLa cells labeling endogenous BLTP2 (green), SCAMP3 (magenta) and GFP-Rab5a (blue), GFP-Rab5a (S34N) or GFP-Rab5a (Q79L) with insets on the right. **E:** Percentage of endogenous BLTP2 on GFP-Rab5a, GFP-Rab5a (S34N) or GFP-Rab5a (Q79L) vesicles from more than three independent experiments as in (**D**). Ordinary one-way ANOVA with Tukey’s multiple comparisons test. Mean ± SD. **F:** Pearson’s correlation coefficient of endogenous BLTP2 vs SCAMP3 in the presence of GFP-Rab5a, GFP-Rab5a (S34N) or GFP-Rab5a (Q79L) from more than three independent experiments as in (**D**). Ordinary one-way ANOVA with Tukey’s multiple comparisons test. Mean ± SD. **G:** Representative images of fixed HeLa cells labeling endogenous BLTP2 (magenta) and GFP-Rab5a (Q79L) upon scrambled or SCAMP3 siRNA with insets on the right. **H:** Pearson’s correlation coefficient of endogenous BLTP2 vs GFP-Rab5a (Q79L) from more than three independent experiments as in (**G**). Two-tailed unpaired Student’s t test. Mean ± SD. Scale bars: 10 μm in the whole cell images and 2 μm in the insets (A, D & G).

While SCAMP3 was found to be located in both endosomes and the Golgi apparatus, BLTP2 was preferentially associated with endosomal SCAMP3 rather than with Golgi SCAMP3 (Fig. 4C), suggesting that there may be a specific interaction between BLTP2 and SCAMP3 in endosomal compartments. Since SCAMP3 interacted with Rab5a in mouse kidney lysate (Fig. 1D), we hypothesized that the recruitment of BLTP2 by SCAMP3 might be regulated by Rab5a. Indeed, IF imaging showed that WT GFP-Rab5, BLTP2, and SCAMP3 puncta were evidently associated with each other (Fig. 4D, top panel). However, the presence of the dominant-negative mutant Rab5a-S34N completely abolished the association between BLTP2 and SCAMP3, resulting in a loss of BLTP2 puncta on Rab5 vesicles (Fig. 4D, middle panel, 4E). Importantly, the presence of the constitutively active mutant Rab5-Q79L caused a striking increase in endosome size^53^ and relatively more colocalization of BLTP2 and SCAMP3 (Fig. 4D, bottom panel, 4F). Depleting SCAMP3 via siRNAs greatly reduced the recruitment of endogenous BLTP2 to the enlarged endosomes induced by Rab5-Q79L (Fig. 4G,H), indicating that SCAMP3 is indispensable for Rab5-regulated recruitment. Collectively, these results suggest that Rab5 regulates BLTP2 recruitment to endosomal compartments via SCAMP3.

### SCAMP3 recruitment of BLTP2 is attenuated by NEDD4-mediated ubiquitination

The NT domain of SCAMP3 contains a Pro-Pro-Ala-Tyr (PPAY) motif that mediates the protein’s interaction with the ubiquitin E3 ligase NEDD4^49^. Indeed, we found that NEDD4-Halo formed puncta that colocalized with mCh-SCAMP3 (Fig. S3A), and in a series of GFP-trap assays, GFP-SCAMP3 was found to interact with either NEDD4-Halo (Fig. S3B) or endogenous NEDD4 (Fig. S3C). Therefore, we examined whether the ubiquitination of SCAMP3 by NEDD4 affects its interaction with BLTP2 by performing GFP-trap assays, and the results showed that depleting NEDD4 enhanced the BLTP2– SCAMP3 interaction (Fig. 5A). The efficiency and specificity of the siRNA-mediated depletion of NEDD4 were assessed via immunoblotting (Fig. 5B). We also found that SCAMP3-Y53A, a mutant defective in NEDD4 binding, increased the BLTP2–SCAMP3 interaction (Fig. 5C), which suggested that NEDD4 negatively regulated the BLTP2–SCAMP3 interaction via SCAMP3 ubiquitination.

**Fig. 5.**
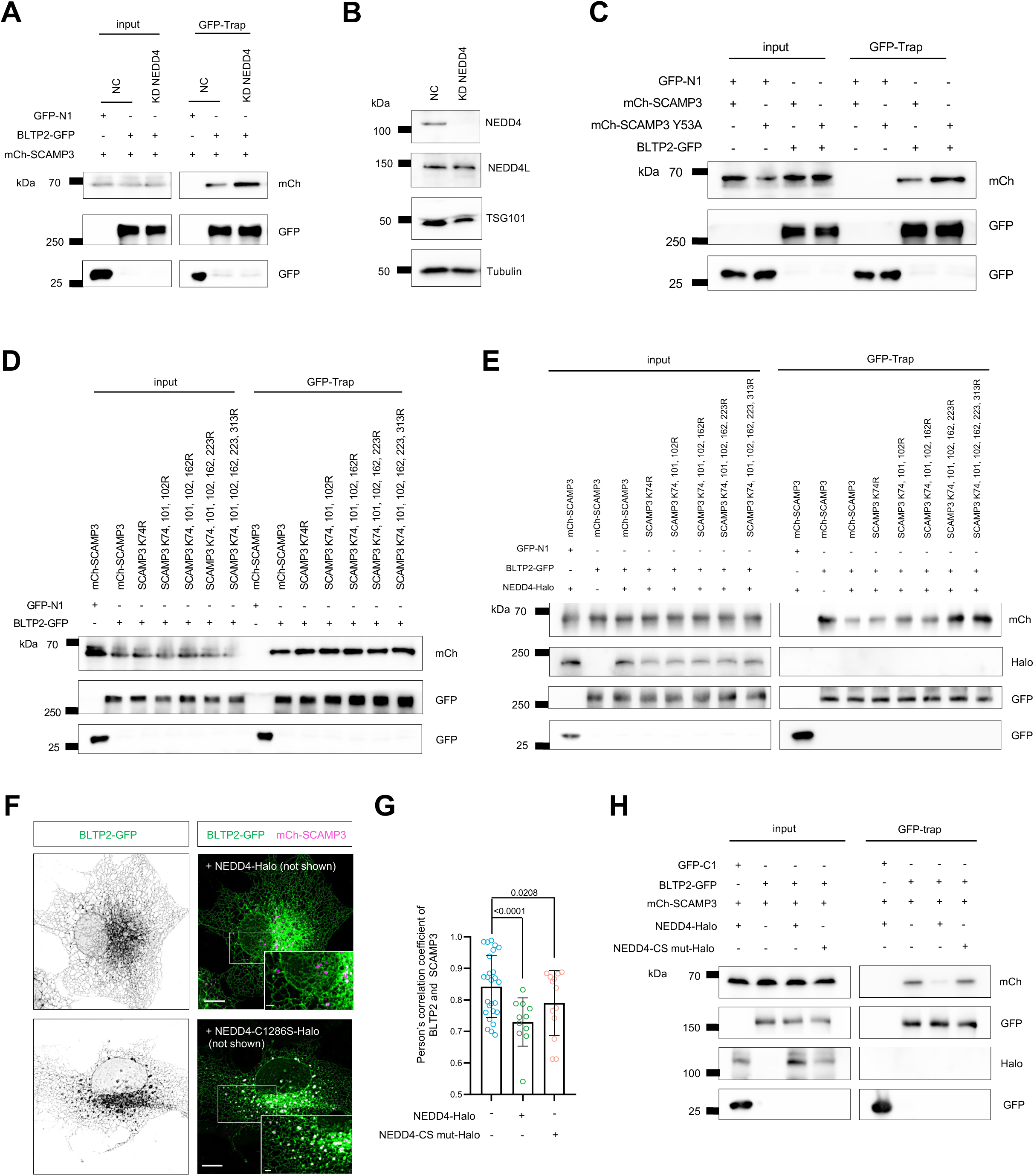
SCAMP3 ubiqutination by NEDD4 attenuates the recruitment of BLTP2. **A:** GFP-Trap assays demonstrate interactions between BLTP2-GFP and mCh-SCAMP3 upon scrambled or NEDD4 siRNA in HEK293 cells. **B:** Immunobloting showing the efficiency of NEDD4 depletion via siRNAs in HEK293 cells as in (**A**). **C:** GFP-Trap assays demonstrate interactions between BLTP2-GFP and mCh-SCAMP3 or mCh-SCAMP3-Y53A in HEK293 cells. **D:** GFP-Trap assays demonstrate interactions between mCh-SCAMP3 mutants and BLTP2-GFP in HEK293 cells. **E:** GFP-Trap assays demonstrate interactions between mCh-SCAMP3 mutants and BLTP2-GFP upon the expression of NEDD4-Halo in HEK293 cells. **F:** Representative images of live COS7 cells expressing BLTP2-GFP (green), mCh-SCAMP3 (magenta) and Halo-NEDD4 or Halo-NEDD4 mutation (C1286S) with insets. **G:** Pearson’s correlation coefficient of BLTP2 vs SCAMP3 upon Halo-NEDD4 or Halo-NEDD4 mutation (C1286S) in more than three independent experiments. Ordinary one-way ANOVA with Tukey’s multiple comparisons test. Mean ± SD. **H:** GFP-Trap assays demonstrate interactions between mCh-SCAMP3 and BLTP2-GFP upon the expression of NEDD4-Halo or NEDD4-C1826S-Halo in HEK293 cells Scale bars: 10 μm in the whole cell images and 2 μm in the insets in (F).

Next, we prepared a series of SCAMP3 mutants that could not be fully ubiquitinated by replacing Lys residues with Arg and examined their ability to interact with BLTP2. We found that the interaction was only moderately enhanced when the K74R mutant was present and that changing additional residues did not further increase the interaction (Fig. 5D). We reasoned that the moderate change observed might have been due to low NEDD4 expression or activity in the HeLa cells. Indeed, the strength of BLTP2–SCAMP3 interaction was considerably reduced when NEDD4 was overexpressed, and the interaction was gradually restored as the number of changed residues increased (Fig. 5E), suggesting that SCAMP3 ubiquitination mediated by NEDD4 attenuates the BLTP2–SCAMP3 interaction.

Accordingly, BLTP2 recruitment by SCAMP3 was significantly hindered by NEDD4-Halo (Fig. 5F, top panel) but not by the E3 ligase-dead mutant NEDD4-C1286S (Fig. 5F, bottom panel), as shown by the colocalization analyses performed using live-cell microscopy (Fig. 5G). In addition, the results of co-IP assays confirmed that the BLTP2–SCAMP3 interaction was considerably reduced when NEDD4-Halo was present compared to the empty vector control, whereas NEDD4-C1286S did not substantially affect the BLTP2–SCAMP3 interaction (Fig. 5H). Together, these results suggest that the ubiquitination of SCAMP3 by NEDD4 negatively regulates the BLTP2–SCAMP3 interaction.

### BLTP2 and SCAMP3 associate with ESCRT machinery

SCAMP3 contains a Pro–Ser–Ala–Pro (PSAP) motif in its Pro-rich region that mediates binding to the ESCRT-II protein TSG101^49^. Accordingly, our results showed that SCAMP3 interacted with TSG101 under physiological conditions (Fig. 1I). Therefore, we investigated whether the BLTP2–SCAMP3 complex associates with TSG101. Live-cell imaging showed that Halo-TSG101 colocalized with BLTP2-GFP and mCh-SCAMP3 at ER–MVB MCSs (Fig. S3D, top panel). Consistently, both the CT of BLTP2 and TSG101 were recruited to mCh-SCAMP3-positive vesicles, suggesting that BLTP2 and TSG101 bind to distinct regions of SCAMP3 (Fig. S3D, middle panel). We next investigated whether other ESCRT proteins associate with BLTP2 and SCAMP3. Vps4B-E235Q, the ATPase-dead mutant of Vps4B, was found to colocalize with BLTP2 and SCAMP3 (Fig. S3D, bottom panel), which suggested that Vps4B associates with the BLTP2– SCAMP3 complex at ER–MVB MCSs. We confirmed the associations between the ESCRT proteins and endogenous, untagged BLTP2 and SCAMP3 via IF and colocalization analyses (Fig. S3F, G). In addition, siRNA-mediated depletion of ESCRT proteins, including TSG101, Vps4B, and HGS, had no effect on the recruitment of BLTP2 by SCAMP3 (Fig. S4H, I).

Thus, our results suggest that there is interplay between the BLTP2–SCAMP3 complex and ESCRTs, and that the recruitment of BLTP2 by SCAMP3 is independent of ESCRTs. They also suggest that the formation of ER–MVB MCSs, which is mediated by the BLTP2–SCAMP3 interaction, may occur upstream of the MVB formation processes that involve ESCRTs.

### BLTP2 is required for ILV/exosome formation during MVB biogenesis

The discovery that BLTP2 is recruited to ER–MVB MCSs by SCAMP3 prompted us to explore its cellular functions. We hypothesized that BLTP2 plays a role in MVB and exosome formation on the basis of three pieces of evidence. First, our co-IP–MS results indicated that BLTP2 may be involved in the formation of exosomes (Fig. S1A), which originate from ILVs in MVBs. Second, BLTP2 and SCAMP3 were found to be spatially associated with ESCRT machinery, the key protein components required for MVB formation. Third, SCAMP3, which we have identified as a BLTP2 adaptor in this study, reportedly promotes the formation of MVBs that may be specialized for exocytosis^11^. Therefore, we sought to determine the role that BLTP2 plays in the formation of MVBs and exosomes.

We began this part of our study by undertaking a proteomic analysis and subsequently identifying SCAMP3 in the exosomal fraction enriched from the culture medium of HeLa cells (Fig. 6A), a result that was consistent with that of a previous study^54^. The purity of the exosomes was assessed via nanoparticle tracking assays (Fig. 6B), transmission electron microscopy (TEM) (Fig. 6C, D), and immunoblotting (Fig. 6E). The results suggested that SCAMP3, like other tetraspanins, was present in MVBs that could facilitate ILV formation and/or the incorporation of specific types of cargo into ILVs. Importantly, the immunoblotting results showed that the number of exosomes in the culture medium was significantly lower in two independent BLTP2-KO clones (KO-5 and KO-7) compared to control cells, with TSG101, CD81, CD9, SCAMP3, and EGFR used as exosomal markers (Fig. 6E). The efficiency of BLTP2 depletion in these two clones were confirmed using dot blots (Fig. S2G, bottom panel).

**Fig. 6.**
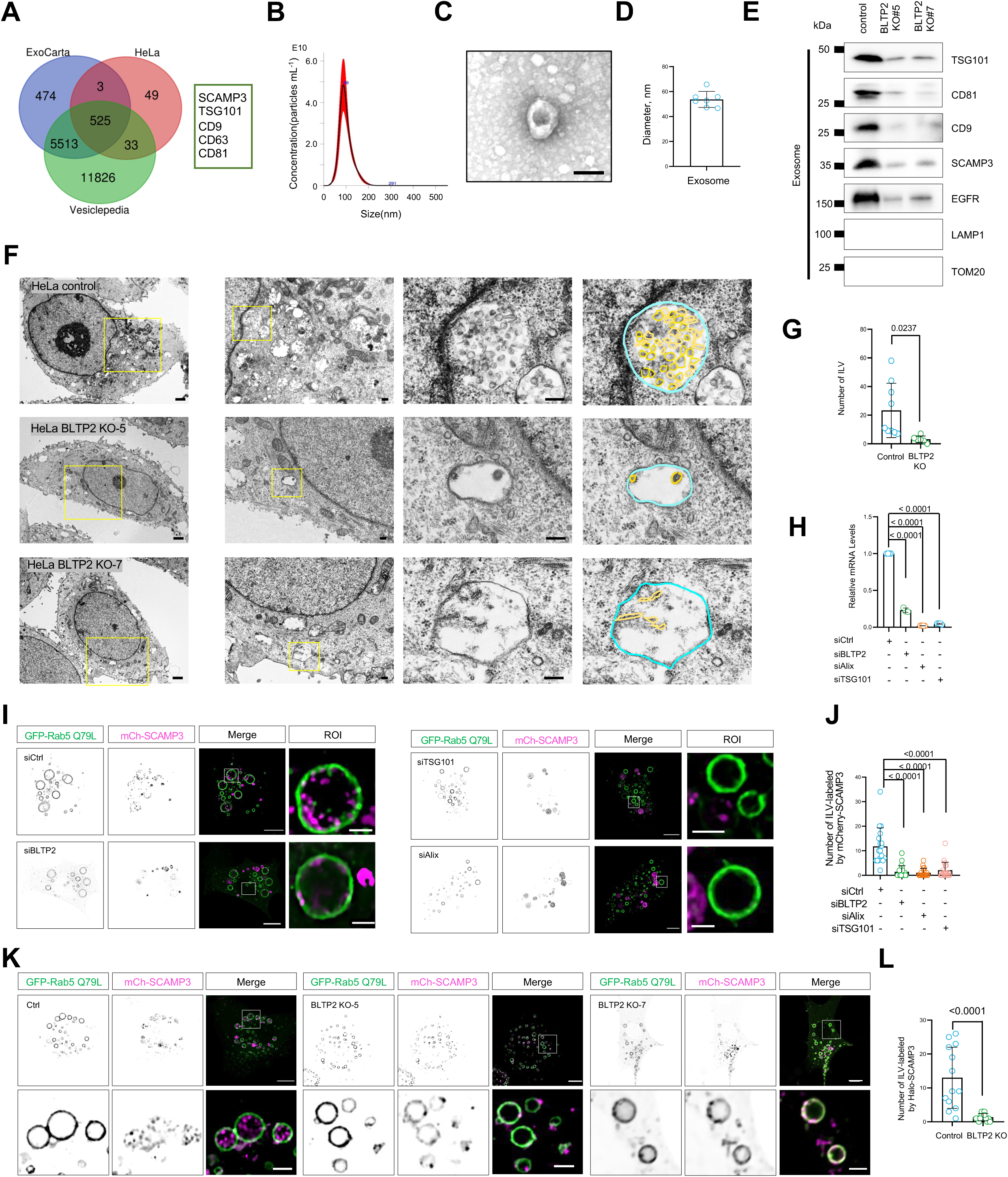
BLTP2 depletion impairs the formation of exosomes/ILVs. **A:** A Venn diagram showing the overlap between the exosomal proteins from the ExoCart database, the Vesiclepedia database and our results from HeLa cells. Some representative overlapped proteins are listed on the right. **B:** Representative NTA analysis on isolated exosomes in this study. The X axis represents the diameters of the vesicles, and the Y axis represents the concentration (particles mL^-1^) of the vesicles. **C:** Representative TEM micrographs showing an example of an exosome isolated in this study. **D:** Diameter of isolated exosomes in TEM as in (**C**) from three independent assays. Mean ± SD. **E:** Immunoblotting of exosome markers (Tsg101, CD81, CD9, SCAMP3) in exosomes purified from control or two BLTP2 KO clones. **F:** Representative TEM micrographs showing examples of MVB in control or two BLTP2 KO clones with insets on the right. **G:** The number of ILVs in control or HeLa BLTP2 KO in three independent assays as in (**F**). Two-tailed unpaired Student’s t test. Mean ± SD. **H:** Relative mRNA Levels in HeLa cells treated with scrambled, BLTP2, TSG101 or Alix siRNAs in three independent experiments. Ordinary one-way ANOVA with Tukey’s multiple comparisons test. Mean ± SD. **I:** Representative images of live HeLa cells expressing GFP-Rab5(Q79L) (green) and mCh-SCAMP3 (magenta) upon scrambled, BLTP2, TSG101 or Alix siRNAs with insets on the right. **J:** The number of ILVs labeled by mCh-SCAMP3 upon scrambled, BLTP2, TSG101 or Alix siRNAs in more than three independent experiments. Ordinary one-way ANOVA with Tukey’s multiple comparisons test. Mean ± SD. **K:** Representative images of live control, BLTP2 KO-5 or BLTP2 KO-7 HeLa cells expressing GFP-Rab5(Q79L, green) and mCh-SCAMP3 (magenta) with insets on the right. **L:** The number of ILVs labeled by mCh-SCAMP3 in control or BLTP2 KO cells in more than three independent experiments. Two-tailed unpaired Student’s t test. Mean ± SD. Scale bars: 10 μm in the whole cell images and 2 μm in the insets (F, I & K).

We hypothesized that the lack of exosomes in the culture medium of the BLTP2-KO cells was due to a defect in ILV formation during MVB biogenesis. Therefore, we performed TEM to directly examine the ILVs. The TEM images showed that there were significantly fewer ILVs in the two BLTP2-KO clones than in the control cells (Fig. 6F, G). These results suggest that BLTP2 is required for efficient ILV/exosome formation.

To confirm the role that BLTP2 plays in ILV formation, we examined ILVs using live-cell imaging. For this purpose, we transfected HeLa cells with the constitutively active Rab5 mutant (Rab5-Q79L) to produce enlarged endosomes and make the ILVs more visible when examined using confocal microscopy. The ESCRT-related proteins ALIX and TSG101 were used as controls. In accordance with previous studies, siRNA-mediated suppression of these two ESCRT proteins significantly reduced the number of ILVs marked by mCh-SCAMP3 compared to the number in the control cells (Fig. 6H, I). This confirmed the key roles that these two proteins play in ILV formation^20, 55^. Importantly, using two different siRNAs to silence BLTP2 greatly reduced the number of endosomal ILVs marked by mCh-SCAMP3 in this experimental system (Fig. 6H, I). We then analyzed ILV formation in BLTP2-KO cells. While multiple mCh-SCAMP3-labeled ILVs were present in the endosomes of the control cells (Fig. 6K, left panel), ILVs were barely present in the endosomes of two BLTP2-KO clones (Fig. 6K, middle and right panels, 6L). Therefore, our results indicate that BLTP2 is required for the formation of ILVs and exosomes.

Given that not all ILVs within MVBs are destined to participate in exocytosis and form exosomes, we undertook a series of experiments to determine whether BLTP2 plays a role in the MVB-mediated degradation pathway. For this, we focused on EGFR, a transmembrane cargo primarily destined for degradation in lysosomes. Knocking out BLTP2 did not completely block the degradation of EGFR; however, it significantly slowed its degradation (Fig. S4A, B). This result suggests that BLTP2 is not required for EGFR degradation but may contribute to the efficient disposal of EGFR via exosomes (Fig. 6D).

### Lipid transfer mediated by BLTP2 is required for ILV/exosome formation

Next, we explored how BLTP2 regulates ILV/exosome formation. As BLTP2 belongs to the BLTP family, we investigated whether ILV/exosome formation is dependent on its lipid transfer activity and, if so, to what extent. We constructed a lipid transfer-deficient BLTP2 mutant (BLTP2-mut), in which several hydrophobic residues in the middle of the hydrophobic groove were changed to charged residues (Fig. S4C) to impede the movement of lipids through the groove^56^, and tested its ability to rescue exosome formation in BLTP2-KO cells. We found that introducing BLTP2-GFP into BLTP2-KO cells efficiently restored the phenotype (Fig. S4D), whereas BLTP2-mut, the lipid transfer-deficient mutant, failed (Fig. S4D). The striking difference between the results obtained with the WT protein and the lipid transfer-deficient mutant was not due to their expression levels, as our immunoblot analysis showed that the proteins were present at similar levels (Fig. S4D). These results suggest that the role BLTP2 plays in ILV/exosome biogenesis depends on its lipid transfer activity.

### BLTP2 KO reduces the levels of PG, BMP/LBPA and PE in MVBs

Given that our results indicated that BLTP2-mediated lipid transfer is critical for ILV/exosome formation, we sought to identify the lipid species associated with BLTP2 in cells. To achieve this, we utilized co-IP followed by non-targeted lipidomic analysis using BLTP2-GFP as the bait^57^. We found that BLTP2-GFP was mainly associated with glycerophospholipids, including phosphatidylcholine (PC), phosphatidylethanamine (PE), phosphoinositide (PI), and PG/BMP (Fig. S5A), and hence showed similar specificity to other Vps13-like proteins^44, 58^. BLTP2-GFP was also associated with sphingolipids (sphingomyelin and ceramide), diglycerides (DG), and triglycerides (TG) (Fig. S5A). However, it should be noted that the associations between BLTP2 and sphingolipids or TG are likely indirect. Interestingly, BLTP2-GFP associated with PG, the precursor of BMP/LBPA, an MVB/late endosome-specific glycerophospholipid (Fig. S5A) essential for ILV formation and late endosome/lysosome functions^20^, although there is relatively little PG in the membranes of HEK293 cells (Fig. S5B). These results suggest that BLTP2 specifically binds to PG.

It has been reported that cone-shaped lipids, including PE, DG, ceramide, cholesterol, and BMP/LBPA, induce negative membrane curvature during ILV formation^6, 21^. Therefore, it is possible that BLTP2 transfers these lipids to facilitate membrane invagination during ILV biogenesis. To test this hypothesis, we isolated endosomal and ER membrane fractions from control and BLTP2-KO cells (Fig. 7A–C) and analyzed the levels of PE, PC, PG, and BMP/LBPA in the fractions using targeted lipidomics. When we examined the endosomal fractions, we found that there was significantly less PG, BMP/LBPA and PE in the BLTP2-KO cell fractions compared to the control cell fractions (Fig. 7D, 7L and 7M) and that the PG and BMP/LBPA levels were reduced to a greater extent than the PE level (Fig. 7E–G). In contrast, the levels of PC, PA, phosphatidylserine (PS), and PI were not substantially affected by knocking out BLTP2 (Figs. 7H–K, and S5C–F). When we examined the ER membrane fractions, we found that the levels of PE and PG were also lower in the BLTP2-KO cell fractions compared to the control cell fractions (Fig. 7D–F), which suggested that knocking out BLTP2 may have affected lipid homeostasis in the ER.

**Fig. 7.**
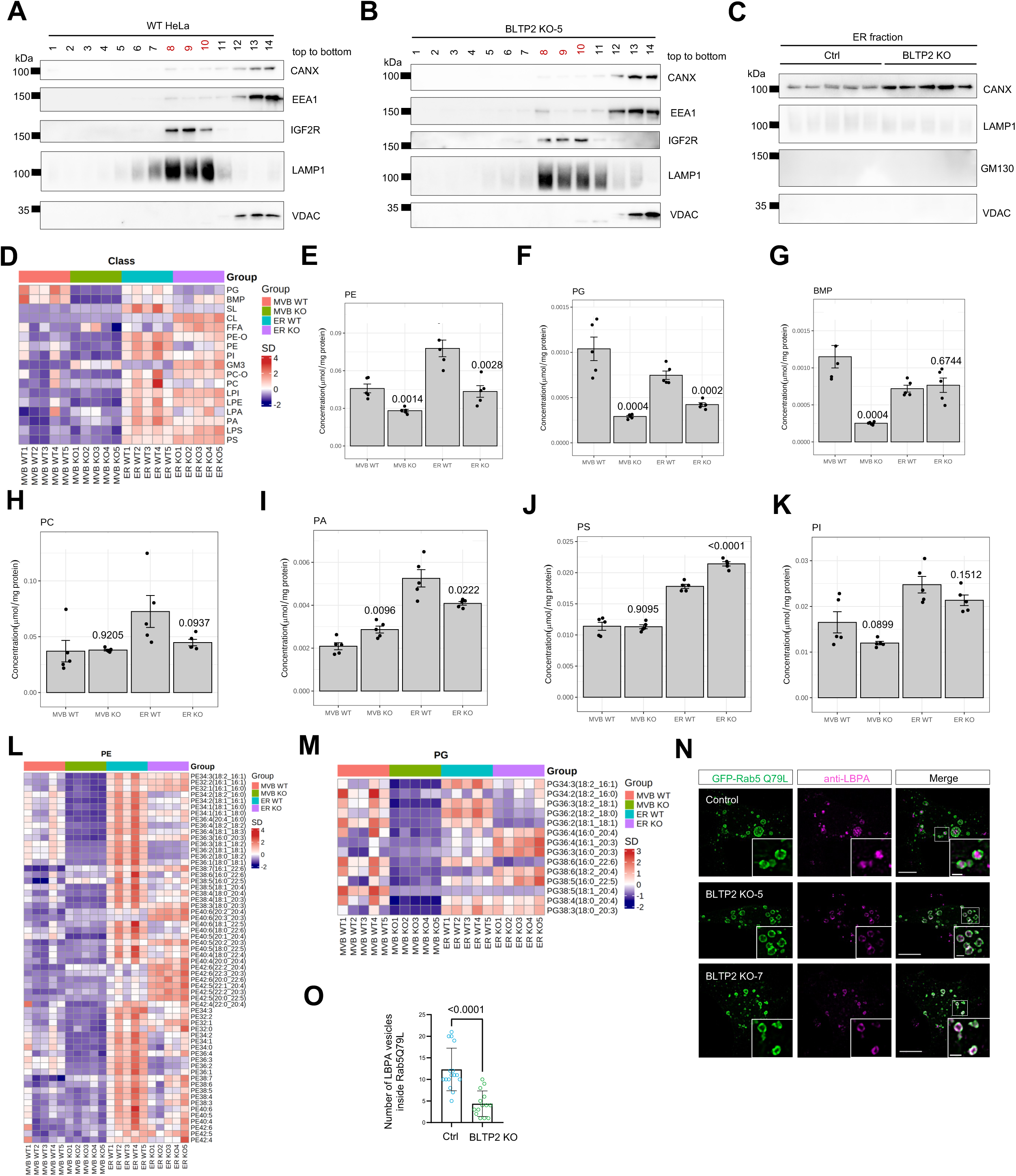
BLTP2 depletion reduced the levels of PE, PG and LBPA/BMP in endosomes. **A, B:** OptiPrep flotation assays showing the distribution of MVB membranes in consecutive 14 fractions from control (**A**) or BLTP2 KO-5 (**B**) HeLa cells. **C:** Western blots of the enriched ER (anti-Calnexin) from control or BLTP2 KO-5 HeLa cells. **D:** Heatmap showing the levels of different lipid species in the endosomal fraction or the ER fraction from control or BLTP2 KO-5 HeLa cells as in (**A-C**) in five independent experiments. **E-K:** The levels of PE (**E**), PG (**F**), LBPA/BMP (**G**), PC (**H**), PA (**I**), PS (**J**) and PI (**K**) in the endosomal fraction or the ER fraction from control or BLTP2 KO-5 HeLa cells as in (**A-C**) in five independent experiments. Two-tailed unpaired Student’s t test. Mean ± SD. **L, M:** Heatmap showing the levels of different PE (**L**) or PG (**M**) species in the endosomal fraction or the ER fraction from control or BLTP2 KO-5 HeLa cells as in (**A-C**) in five independent experiments. **N:** Representative images of fixed control, BLTP2 KO-5 or BLTP2 KO-7 expressing GFP-Rab5(Q79L;green) stained with antibody against LBPA (magenta) with insets. **O:** The number of LBPA vesicles inside Rab5(Q79L) endosomes in control or BLTP2 KO cells in more than three independent experiments. Two-tailed unpaired Student’s t test. Mean ± SD. Scale bars: 10 μm in the whole cell images and 2 μm in the insets in (N).

To further investigate the effect of BLTP2 on BMP/LBPA, we performed IF staining using a monoclonal antibody against BMP/LBPA (6C4^20^) in cells with and without BLTP2. In contrast to the control cells (transfected with the Rab5-Q79L mutant), which exhibited several BMP/LBPA puncta in the endosomal lumen, BMP/LBPA puncta were almost completely absent from the endosomal lumen in the two tested BLTP2-KO clones; some BMP/LBPA remnants were observed in the limiting membranes of the endosomes (Fig. 7O, P). To examine the acute rather than chronic effects of BLTP2 deficiency on BMP/LBPA, we depleted BLTP2 by two different siRNAs. The phenotype was also observed in BLTP2-depleted HeLa cells (Fig. S5G, H). Collectively, these results indicate that BLTP2 plays a specific role in the regulation of the levels of PG, BMP/LBPA, and PE in endosomes, which may contribute to ILV formation.

### BLTP2 promotes cell proliferation and tumorigenicity via exosomes

We found that knocking out BLTP2 results in embryonic lethality in mice (unpublished data), indicating that BLTP2 is essential for life, in line with a recent study^47^. In this study, we performed a series of cell proliferation assays to assess the effect of BLTP2 on the proliferation of HeLa cells. The results showed that knocking out BLTP2 significantly impaired the proliferation of HeLa cells and that adding purified exosomes from the culture medium of WT cells significantly restored cell proliferation in a dose-dependent manner (Fig. 8A). This indicated that BLTP2-mediated exosome formation contributes to cell proliferation.

**Fig. 8.**
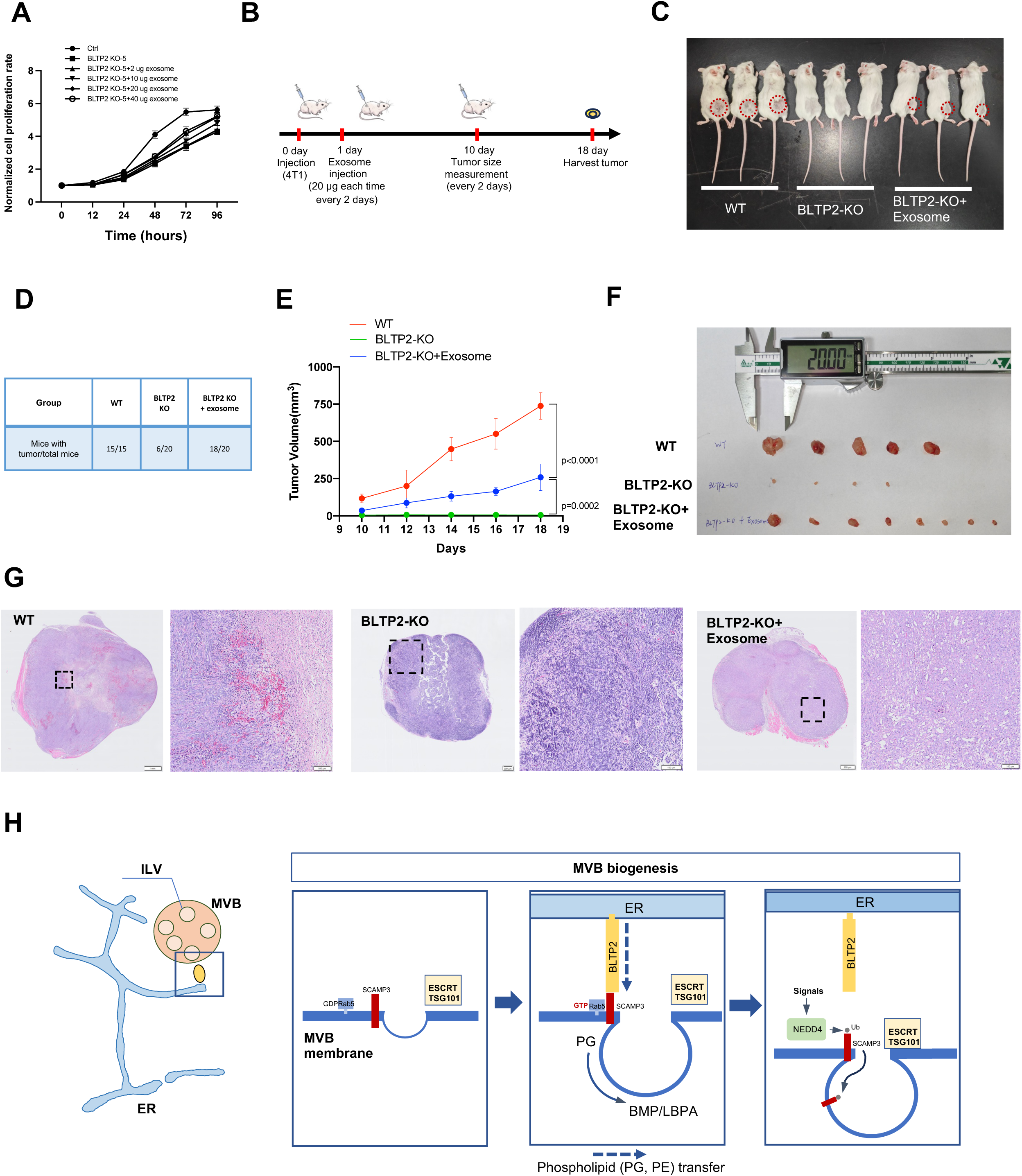
BLTP2 promotes tumor growth dependent of the exosome pathway. **A:** Normalized luminescence of HeLa control, BLTP2 KO-5 or BLTP2 KO-5 administered with different amounts of exosomes isolated from WT HeLa cells in 3 independent experiments. Mean ± SD. **B:**Schematic diagram of the xenograft assay using mice administered with exosomes purified from the culture medium of WT 4T1 cells. **C:** Representative mice in the xenograft assay as in **(B)** 18 days after injecting 4T1 cells. **D:** The chance of tumor development in mice that received WT (15/15, 100%), BLTP2 depleted (6/20, 33.3%) and BLTP2-depleted cell administered with exosomes (18/20, 90%). The quantification was based on three independent assasys. **E:** The measurement of tumor size at different time points as in **(B)** from three independent assays. Two-tailed unpaired Student’s t test. Mean ± SD. **F:** Representative images of harvested tumors. **G:** Hematoxylin and eosin staining of histological sections of tumors as shown in **(F)**. Representative of eight images per group; 12-mm sections; **H:** The working model of BLTP2 in the formation of exosomes/ILVs during MVB biogenesis. Scale bar: 1 mm in the whole cell images and 100 μm in the insets in (G, left panel); 200 μm in the whole cell images and 100 μm in the insets in (G, middle panel); 500 μm in the whole cell images and 100 μm in the insets in (G, right panel);

Next, we investigated whether and to what extent the role of BLTP2 in tumorigenesis is dependent on exosome using a 4T1 xenograft model. For this, we implanted control or BLTP2-depleted 4T1 cells subcutaneously into BALB/c mice and evaluated the tumor size. Most (70%) of the mice that received BLTP2-depleted cells could not develop tumors and the remaining 30% of mice developed considerably smaller tumors than those that received control cells (Fig. 8B, C). These results provide further evidence that BLTP2 plays a crucial role in tumor cell survival. It should also be noted that administering purified exosomes from the culture medium of WT cells via subcutaneous injection near the xenograft injection site significantly increased the chance of tumor development in the mice that received BLTP2-KO cells (90% vs 30%; Fig. 8D). In addition, exosome administration significantly increased the size of tumors from the mice that received BLTP2-depleted cells (Fig. 8E, F).

We next performed hematoxylin and eosin (H&E) staining to analyze tumor morphology. Strikingly, tumors derived from BLTP2-KO cells exhibited markedly fewer intratumoral capillaries compared to WT tumors, whereas exosome supplementation significantly restored capillary formation in BLTP2-KO tumors (Fig. 8G). These findings suggest that BLTP2 may regulate tumor angiogenesis through exosome-mediated mechanisms. Together, these results suggest that BLTP2-mediated exosome formation promotes cell proliferation and tumorigenesis.

## Discussion

In this study, we have revealed for the first time that BLTP2 plays an essential role in ILV/exosome formation. The working model developed from our results is shown in Fig. 8H. At the onset of ILV/exosome formation during MVB biogenesis, Rab5 induces the recruitment of ER-anchored BLTP2 to ER–endosome MCSs via the endosome-resident tetraspanin SCAMP3. This recruitment is mediated by electrostatic interactions between an α-helix located at the CT of BLTP2 and an α-helix located at the NT of SCAMP3. At SCAMP3-dependent ER–MVB MCSs, BLTP2 transports the BMP/LBPA precursor PG and PE from the ER to MVBs and promotes BMP/LBPA synthesis and membrane invagination and consequently facilitates ILV/exosome formation. Then, the BLTP2 adaptor SCAMP3 is ubiquitinated by the E3 ligase NEDD4, which inhibits the BLTP2–SCAMP3 interaction and causes the ER to lose contact with MVBs, followed by SCAMP3 invagination.

It is worth noting that yeast Vps13 is recruited to several types of MCSs via different adaptors^59^ and that mammallian VPS13A can be recruited to ER-mitochondrial and ER-plasma membrane (PM) MCSs^58, 60–63^. In addition, Fmp27 and Hob2, the yeast homologues of BLTP2, have been found at ER-PM or ER–mitochondrial MCSs^32, 47^. Furthermore, BLTP2 has been found at WDR44-dependent ER– endosome^46^ and ER-PM MCSs via multiple interactions with phosphoinositides, FAM102A and FAM102B, and also N-BAR domain proteins in mammallian cells^64^. These findings suggest that the localization of Vps13-like proteins, including BLTP2, may be versatile and can be regulated, depending on the functional state of the cell. Therefore, it is plausible that BLTP2 is recruited to multiple types of MCSs under physiological or pathological conditions by different adaptors.

BMP/LBPA is an unconventional phospholipid that is enriched in ILVs, essential for MVB formation, and a potent stimulator of lysosomal function. Alterations in the level of BMP/LBPA have been linked to neurodegeneration, and its accumulation is increasingly recognized as a key response to lysosomal dysfunction^65^. The BMP/LBPA precursor PG is an intermediate lipid in the mitochondrial cardiolipin pathway, and no other studies have determined how PG is transported to MVBs for BMP/LBPA synthesis. Our results imply that BLTP2 transports PG from the ER to MVBs for BMP/LBPA synthesis and that the ER may serve as a depot for PG during its transfer from mitochondria to MVBs.

A lack of BLTP2 leads to embryonic lethality in mice (unpublished results); hence, BLTP2 plays an essential physiological role. In contrast, relatively high levels of BLTP2 are associated with acute monocytic leukemia^66^ and breast cancer^67^. Together, these findings indicate that regulating BLTP2 is fundamental to maintaining health. In this study, we examined the mechanisms underlying BLTP2 regulation and found that Rab5 controls the recruitment of BLTP2 to ER–endosome MCSs and that the ubiquitin E3 ligase activity of NEDD4 contributes to the disassembly of these sites. Thus, these two molecules (Rab5 and NEDD4) play critical roles in BLTP2 regulation and exosome formation, which are essential in both cell survival and tumorigenicity. Overall, our findings provide insight into the mechanistic role that BLTP2 plays in physiological processes and cancer development and indicate that the BLTP2-mediated transfer of PG from the ER to MVBs is a key step in BMP/LBPA synthesis in endosomes.

## Materials and methods

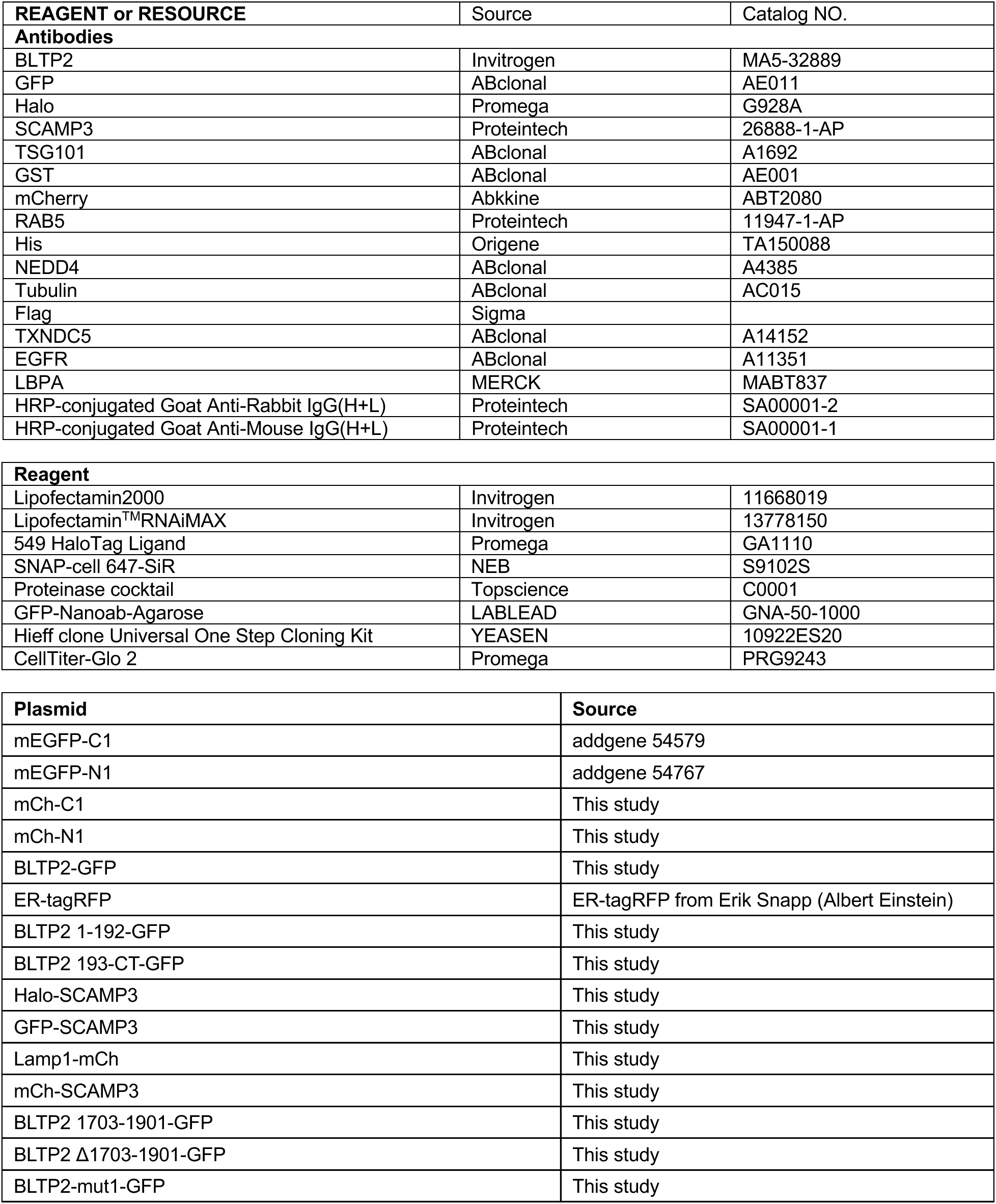
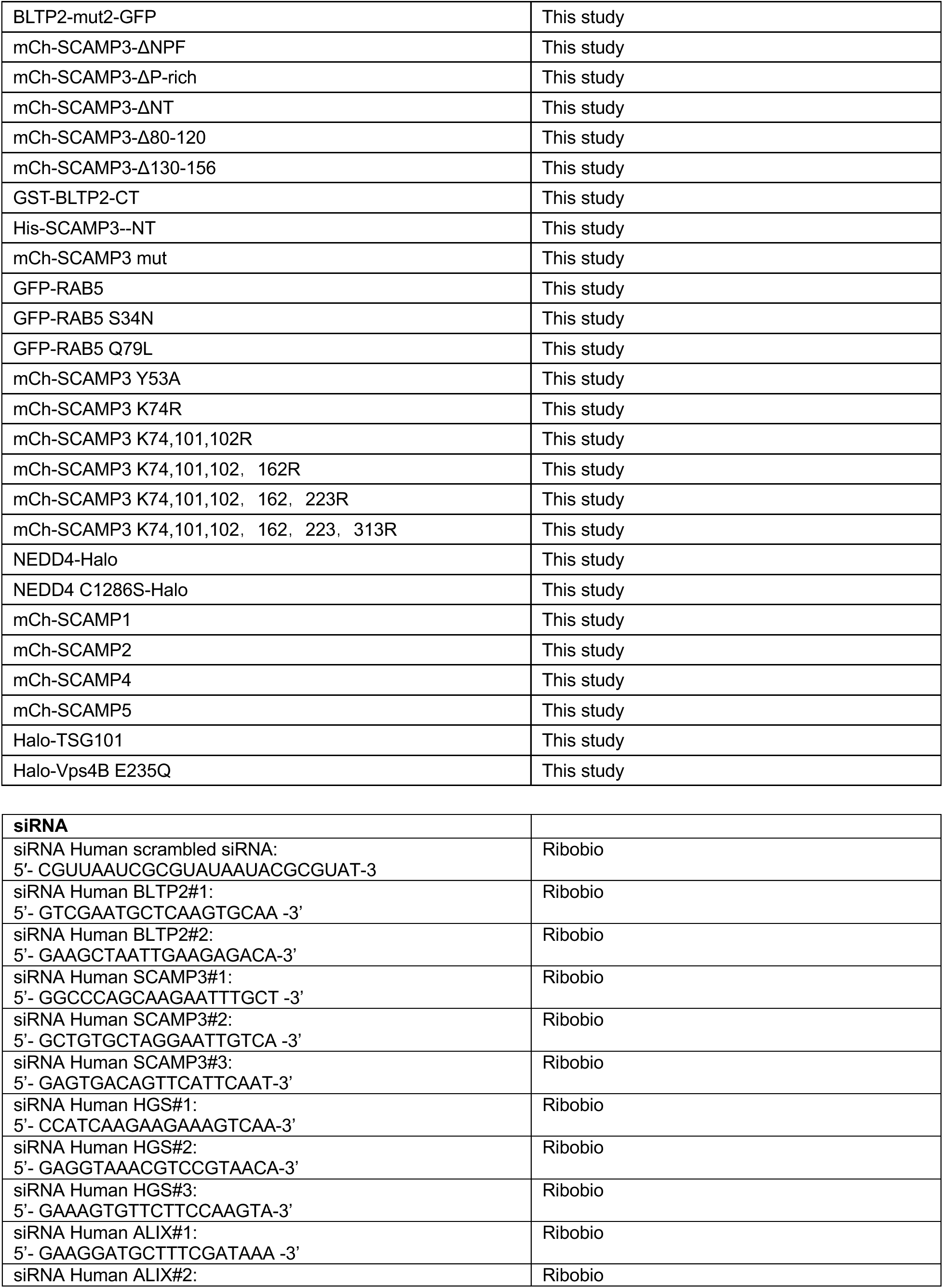
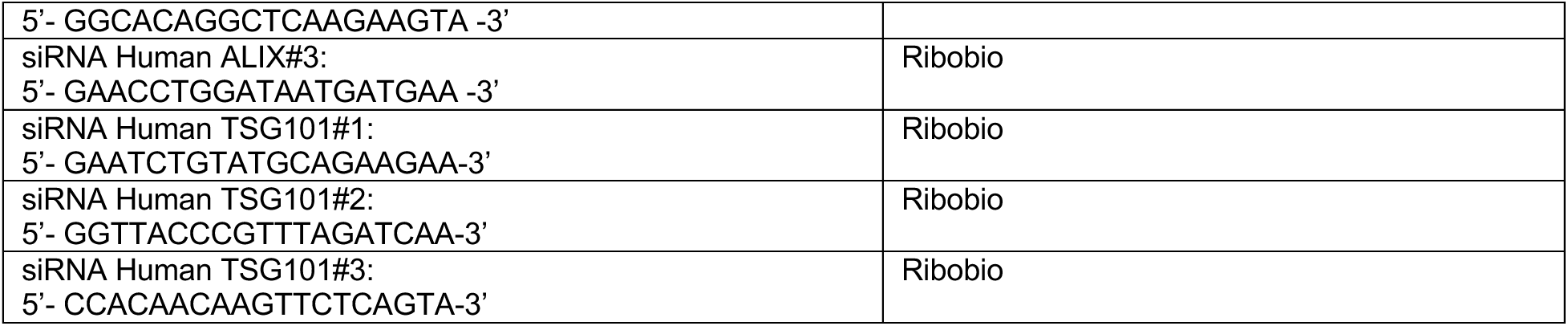

### Cell culture, transfection, RNAi

The African green monkey kidney fibroblast-like COS7 cell line (ATCC), HeLa (ATCC), and human embryonic kidney 293T (ATCC) were grown in DMEM (Invitrogen) supplemented with 10% fetal bovine serum (Gibco) and 1% penicillin/streptomycin. All the cell lines used in this study were confirmed free of mycoplasma contamination.

Transfection of plasmids and RNAi oligos was performed with Lipofectamine 2000 and RNAi MAX, respectively. For transfection, cells were seeded ∼16 h before transfection with 4 x 10^5^ cells per well in a six-well dish. Plasmid transfections were performed in OPTI-MEM (Invitrogen) with 2 μl Lipofectamine 2000 per well for 6 hours. Cells were then trypsinized and replicated onto confocal glass bottom dishes with ∼3.5 x 105 cells per well. Cells were imaged ∼16-24 hours after transfection in live cell medium (DMEM with 10% FBS and 20 mM Hepes without penicillin or streptomycin)–. For all transfection experiments in this study, the following amounts of DNA were used per 3.5 cm well (individually or combined for co-transfection): 1000ng for BLTP2-GFP and its mutants, 300 ng for mCherry-SCAMP3 and its mutants; 500 ng for ER-tagRFP, OFP-Rab5a and other constructs. For siRNA transfections, cells were plated on 3.5 cm dishes at a density of 30–40% and 2 μl Lipofectamine RNAimax (Invitrogen) and 50 ng siRNA per well were used. 48 hours after transfection, a second round of transfection with 50 ng siRNAs was performed. Cells were analyzed for suppression 24 hours after the second transfection.

### BLTP2 KO HeLa cell lines

To generate BLTP2 KO cell lines, two gRNAs (5’-gcttggctgtgctcacctcc -3’ and 5’-cctctctctttcggctcagtct-3’) were used to delete 269 bp of exon 16 of the BLTP2 gene (Fig. S2C, D). The complementary gRNAs were annealed and subcloned into the pSpCas9(BB)-2A-GFP (pX-458) vector (#48138; Addgene) between BbsI endonuclease restriction sites. After transfection, individual cells with GFP fluorescent signals were sorted by FACS in 96-well plates and the growing monoclonal clones were verified by PCR (Fig. S2E). The presence of mutations was confirmed by Sanger sequencing. The mutations resulted in code-shifting and premature termination in the BLTP2 coding sequence, and protein expression was verified by qPCR (Fig. S2F) and dot-blot analysis (Fig. S2G). Two validated KO clones (#5 and #7) were used in this study.

### GFP-trap assay

GFP-trap was used for detection of protein–protein interactions and the GFP-Trap assays were performed according to the manufacturer’s protocol. 5% input was used in GFP traps unless otherwise indicated. Briefly, cells were lysed in Lysis buffer (50 mM Tris-Cl pH 7.5, 150 mM NaCl, 0.5 mM EDTA, 0.5 % NonidetTM P40 Substitute). Lysates were centrifuged at 13,000 rpm for 10 min at 4°C and pellets were removed. Supernatants were incubated with GFP-Trap agarose beads for 1h at 4°C with gentle rocking. After washing four times with wash buffer (50 mM Tris-HCl pH 7.5, 150 mM NaCl, 0.5 mM EDTA, 1x Proteases Inhibitor cocktail), beads were boiled with SDS sample buffer. Proteins of interest were analyzed by immunoblotting. 5% input was used in GFP-traps unless otherwise indicated.

### Endogenous immunoprecipitation

Cells lysates were incubated with 1 μg indicated antibody for 8 h at 4°C with gentle rocking, with IgG as a negative control. Then the mixture was incubated with protein A/G beads for another 12 h. The beads were washed three times with TBS, then boiled with SDS sampling buffer.

### Protein expression and purification

His-tag or GST-tag construct was transformed into TSsetta (DE3) chemically competent cells (TSC04; Tsingke). The cells were incubated at 37°C until the optical density (OD) at 600 nm reached 0.6–0.8. They were then infused overnight at 16°C with 1 mM IPTG. The cells were lysed by sonication. The cell lysates were centrifuged at 14,000 g for 30 minutes. The supernatant was incubated with Ni-NTA resin (G600033-0100, Sangon, for His fusion protein) or GST-Tag resin (C600031-0025, Sangon, for GST fusion protein), then the resins were run in gravity flow.

### GST-pull down assays

The indicated GST proteins were incubated with GST-tag resin at 4 °C for 12 hours, followed by washing 10 times with HNM buffer (20mM HEPES, pH 7.4, 100 mM NaCl, 5 mM MgCl2, 1 mM DTT, 0.2%NP-40) and centrifugation at 1,000 g for 2 minutes. The GST resin was incubated with indicated His fusion proteins at 4 °C for another 12 hours. After washing 10 times with HNM buffer, the beads were boiled with SDS sampling buffer. Western blotting was performed using anti-GST or His antibodies.

### Exosome isolation

Cells were cultured in DMEM (Invitrogen) supplemented with 10% exosome depleted fetal bovine serum (Gibco) for 48 h. The cell culture medium was centrifuged at 2,000g for 30 minutes at 4 °C to remove cell debris and dead cells, followed by another centrifuge at 12,000g for 30 minutes at 4 °C to remove microbubbles or protein aggregates. If there are many impurities (with obvious sediment), centrifuge the supernatant at 12,000g for 30 minutes and collect the supernatant until there is no obvious sediment. The supernatant was filtered with a 0.22 μm filter to remove apoptotic bodies and microbubbles. Collect the cell culture medium in a 50 mL centrifuge tube. Exosomes were isolated from these supernatants using the EXODUS platform according to manufacturer instructions.

### Dot Blot assay

After the protein samples (2 µL containing 10 µg proteins) were spotted onto nitrocellulose membrane, the membrane was placed in a plastic container and sequentially incubated in blocking buffer (5% BSA in TBS-T) 1 h at room temperature, followed by the incubation with primary antibody in TBS-T overnight at 4 °C, and then secondary antibody conjugated to HRP for 30 min at room temperature. The membrane was washed three times with TBS-T (1 x 15 min and 2 x 5 min), then once with TBS (5 min).

### Immunofluorescence staining

Cells were fixed with 4% PFA (paraformaldehyde, Sigma) in PBS for 20 min at room temperature. After washing with PBS three times, cells were permeabilized with 0.5% Triton X-100 in PBS for 10 min on ice. Cells were then washed three times with PBS, blocked with 3% BSA in PBS for 1 h, incubated with primary antibodies in diluted blocking buffer overnight, and washed with PBS three times. Secondary antibodies were applied for 1 h at room temperature. After washing with PBS three times, samples were mounted on Vectashield (H-1000; Vector Laboratories). For BLTP2 staining, cells were fixed with pre-cooled acetone at -20°C for 10 minutes, then blocked with 3% BSA.

### Live imaging by high-resolution confocal microscopy

Cells were grown on 35 mm glass-bottom confocal MatTek dishes and were loaded to a laser scanning confocal microscope (LSM900, Zeiss, Germany) equipped with multiple excitation lasers (405 nm, 458 nm, 488 nm, 514 nm, 561 nm and 633 nm) and a spectral fluorescence GaAsP array detector. Cells were imaged with the 63×1.4 NA iPlan-Apochromat 63 x oil objective using the 405 nm laser for BFP, 488 nm for GFP, 561nm for tagRFP or mCherry and 633nm for Janilia Fluo® 646 HaloTag® ligand.

### Electron microscopy

Wild-type or BLTP2 knockout HeLa cells were fixed with 2.5% glutaraldehyde in 0.1 M Phosphate buffer, pH 7.4 for 2 h at room temperature. After washing three times with 0.1 M phosphate buffer, cells were scraped and collected with 0.1 M PBS followed by centrifugation at 1,100 g. The pellet was resuspended in PBS (0.1 M), and centrifuged at 1,100 g for 10 min. This step was repeated three times. The samples were post-fixed with pre-cold 1% OsO4 in 0.1M PBS for 2-3 hour at 4°C, followed by rinsing with PBS for three times (3 × 20 min). The samples were dehydrated in graded ethanol (50%, 70%, 85%, 90%, 95%, 2x100%) for 15 min in each condition. The penetrations were performed in an order of acetone-epoxy (2:1); acetone-epoxy (1:1); epoxy. Each round of penetration was performed at 37°C for 12 hours. The samples were embedded in epoxy resin using standard protocols^68^. Sections parallel to cellular monolayers were obtained using a Leica EM UC7 with the thickness of 60-100 nm and examined under HT7800/HT7700.

### Cell proliferation assay

Cells were seeded in 96-well plate with a density of 3000 cells per well. After ∼24 hours, the medium were replaced by 100 μL detection medium, which was prepared by 1:1 mixature of a volume of CellTiter-Glo2 Reagent and culture medium in a well of 96-well plate, followed by incubation on an orbital shaker for 12 min at room temperature. The luminescence was recored.

### Mass spectrometry for identification of BLTP2-GFP interacting proteins

The identification of BLTP2-GFP-interacting proteins by MS was described in our previous study^50^. Briefly, the bound proteins were extracted from GFP-Trap agarose beads using SDT lysis buffer (4% SDS, 100 mM DTT, 100 mM Tris-HCl, pH 8.0), followed by sample boiling for 3 min and further ultrasonicated. Undissolved beads were removed by centrifugation at 16,000 g for 15 min. The supernatant, containing proteins, was collected. Protein digestion was performed with the FASP method. Briefly, the detergent, DTT, and IAA in the UA buffer were added to block-reduced cysteine. Finally, the protein suspension was digested with 2 μg trypsin (Promega) overnight at 37°C. The peptide was collected by centrifugation at 16,000 g for 15 min. The peptide was desalted with C18 StageTip for further LC-MS analysis. LC-MS/MS experiments were performed on a Q Exactive Plus mass spectrometer that was coupled to an Easy nLC (Thermo Fisher Scientific). Peptide was first loaded to a trap column (100 µm x 20 mm, 5 µm, C18, Dr Maisch GmbH, Ammerbuch, Germany) in buffer A (0.1% formic acid in water). Reverse-phase high-performance liquid chromatography (RP-HPLC) separation was performed using a self-packed column (75 µm x 150 mm; 3 µm ReproSil-Pur C18 beads, 120 Å, Dr Maisch GmbH, Ammerbuch, Germany) at a flow rate of 300 nl/min. The RP-HPLC mobile phase A was 0.1% formic acid in water, and B was 0.1% formic acid in 95% acetonitrile. The gradient was set as following: 2%–4% buffer B from 0 min to 2 min, 4% to 30% buffer B from 2 min to 47 min, 30% to 45% buffer B from 47 min to 52 min, 45% to 90% buffer B from 52 min and to 54 min, and 90% buffer B kept until to 60 min. MS data was acquired using a data-dependent top20 method dynamically choosing the most abundant precursor ions from the survey scan (350– 1800 m/z) for HCD fragmentation. A lock mass of 445.120025 Da was used as internal standard for mass calibration. The full MS scans were acquired at a resolution of 70,000 at m/z 200, and 17,500 at m/z 200 for MS/MS scan. The maximum injection time was set to 50 ms for MS and 50 ms for MS/ MS. Normalized collision energy was 27 and the isolation window was set to 1.6 Th. Dynamic exclusion duration was 60 s. The MS data were analyzed using MaxQuant software version 1.6.1.0. MS data were searched against the UniProtKB Human norvegicus database (36,080 total entries, downloaded 08/14/2018). Trypsin was seleted as the digestion enzyme. A maximum of two missed cleavage sites and the mass tolerance of 4.5 ppm for precursor ions and 20 ppm for fragment ions were defined for database search. Carbamidomethylation of cysteines was defined as a fixed modification, while acetylation of protein N-terminal, oxidation of Methionine were set as variable modifications for database searching. The database search results were filtered and exported with a <1% false discovery rate (FDR) at peptide-spectrum-matched level, and protein level, respectively.

### Non-targeted lipidomics using LC-MS/MS

The identification of BLTP2-GFP-assocaiting lipids in HEK293 cells was described in our previous study^50^. Briefly, to extract lipids, 1 ml methyl tert-butyl ether (MTBE) was added to GFP-Trap agarose beads and the samples were shaken for 1 h at room temperature. Next, phase separation was induced by adding 250 µL water, letting it sit for 10 min at room temperature and centrifuging for 15 min at 14,000 g, 4°C. Because of the low density and high hydrophobicity of MTBE, lipids and lipophilic metabolites are mainly extracted to the upper MTBE-rich phase. The lipid was transferred to fresh tubes and dried with nitrogen. Additionally, to ensure data quality for metabolic profiling, quality control (QC) samples were prepared by pooling aliquots from representative samples for all of the analysis samples, and were used for data normalization. QC samples were prepared and analyzed with the same procedure as that for the experiment samples in each batch. Dried extracts were then dissolved in 50% acetonitrile. Each sample was filtered with a disposable 0.22 μm cellulose acetate and transferred into 2 ml HPLC vials and stored at -80°C until analysis. For UHPLC-MS/MS analysis, lipid analysis was performed on Q Exactive orbitrap mass spectrometer (Thermo Fisher Scientific) coupled to a UHPLC system Ultimate 3000 (Thermo Fisher Scientific). Samples were separated using a Hypersil GOLD C18 column (100 x 2.1 mm, 1.9 µm) (Thermo Fisher Scientific). Mobile phase A was prepared by dissolving 0.77 g of ammonium acetate to 400 ml of HPLC-grade water, followed by adding 600 ml of HPLC-grade acetonitrile. Mobile phase B was prepared by mixing 100 ml of acetonitrile with 900 ml isopropanol. The flow rate was set as 0.3 mL/min. The gradient was 30% B for 0.5 min and was linearly increased to 100% in 10.5 min, and then maintained at 100% in 2 min, and then reduced to 30% in 0.1 min, with 4.5 min re-equilibration period employed. Both electrospray ionization (ESI) positive-mode and negative-mode were applied for MS data acquisition. The positive mode of spray voltage was 3.0 kV and the negative mode 2.5 kV. The ESI source conditions were set as follows: heater temperature of 300°C, Sheath Gas Flow rate, 45arb, Aux Gas Flow Rate, 15 arb, Sweep Gas Flow Rate, 1arb, Capillary Temp, 350°C, S-Lens RF Level, 50%. The full MS scans were acquired at a resolution of 70,000 at m/z 200, and 17,500 at m/z 200 for MS/MS scans. The maximum injection time was set to for 50 ms for MS and 50 ms for MS/MS. MS data was acquired using a data-dependent Top10 method dynamically choosing the most abundant precursor ions from the survey scan (200–1500m/z) for HCD fragmentation. Stepped normalized collision energy was set as 15, 25, 35 and the isolation window was set to 1.6 Th. QC samples were prepared by pooling aliquots that were representative of all samples under analysis, and used for data normalization. Blank samples (75% acetonitrile in water) and QC samples were injected every six samples during acquisition.

For data preprocessing and filtering, lipids were identified and quantified using LipidSearch 4.1.30 (Thermo, CA). Mass tolerance of 5 ppm and 10 ppm were applied for precursor and product ions. Retention time shift of 0.25 min was performed in ‘alignment’. M-score and chromatographic areas were used to reduce false positives. The lipids with less than 30% relative standard deviation (RSD) of MS peak area in the QC samples were kept for further data analysis. SIMCAP software (Version 14.0, Umetrics, Sweden) was used for all multivariate data analyses and modeling. Data were meancentered using Pareto scaling. Models were built on principal component analysis (PCA), orthogonal partial least-square discriminant analysis (PLS-DA) and partial least-square discriminant analysis (OPLS-DA). All the models evaluated were tested for over fitting with methods of permutation tests. The descriptive performance of the models was determined by R2X (cumulative) [perfect model: R2X (cum)=1] and R2Y (cumulative) [perfect model: R2Y (cum)=1] values while their prediction performance was measured by Q2 (cumulative) [perfect model: Q2 (cum)=1] and a permutation test (n=200). The permuted model should not be able to predict classes – R2 and Q2 values at the Y-axis intercept must be lower than those of Q2 and the R2 of the non-permuted model. OPLS-DA allowed the determination of discriminating metabolites using the variable importance on projection (VIP). The VIP score value indicates the contribution of a variable to the discrimination between all the classes of samples. Mathematically, these scores are calculated for each variable as a weighted sum of squares of PLS weights. The mean VIP value is 1, and usually VIP values over 1 are considered as significant. A high score is in agreement with a strong discriminatory ability and thus constitutes a criterion for the selection of biomarkers. The discriminating metabolites were obtained using a statistically significant threshold of variable influence on projection (VIP) values obtained from the OPLS-DA model and two-tailed Student’s t-test (P-value) on the normalized raw data at univariate analysis level. The P-value was calculated by one-way analysis of variance (ANOVA) for multiple groups analysis. Metabolites with VIP values greater than 1.0 and P-value less than 0.05 were considered to be statistically significant metabolites. Fold change was calculated as the logarithm of the average mass response (area) ratio between two arbitrary classes. On the other side, the identified differential metabolites were used to perform cluster analyses with R package.

### Targeted lipidomics using LC-MS/MS

#### Lipid extraction

Lipids were extracted from approximately one million cells using a modified version of the Bligh and Dyer’s method as described previously^69^. Briefly, cells were homogenized in 750 µL of chloroform: methanol: MilliQ H_2_O (3:6:1) (v/v/v). The homogenate was then incubated at 1500 rpm for 1h at 4℃. At the end of the incubation, 350 µL of deionized water and 250 µL of chloroform were added to induce phase separation. The samples were then centrifuged and the lower organic phase containing lipids was extracted into a clean tube. Lipid extraction was repeated once by adding 450 µL of chloroform to the remaining cells in aqueous phase, and the lipid extracts were pooled into a single tube and dried in the SpeedVac under OH mode. Samples were stored at -80℃ until further analysis. Upper aqueous phase and cell pellet were dried in a SpeedVac under H_2_O mode. Total protein content was determined from the dried pellet using the Pierce® BCA Protein Assay Kit according to the manufacturer’s protocol.

#### Lipidomics analyses

Lipidomic analyses were conducted at LipidALL Technologies using a ExionLC-AD coupled with Sciex QTRAP 6500 PLUS as reported previously ^70^. Separation of individual lipid classes of polar lipids by normal phase (NP)-HPLC was carried out using a TUP-HB silica column (i.d. 150x2.1 mm, 3 µm) with the following conditions: mobile phase A (chloroform: methanol:ammonium hydroxide, 89.5:10:0.5) and mobile phase B (chloroform:methanol:ammonium hydroxide:water, 55:39:0.5:5.5). MRM transitions were set up for comparative analysis of various polar lipids. Individual lipid species were quantified by referencing to spiked internal standards. d_9_-PC32:0(16:0/16:0), d_9_-PC36:1p(18:0p/18:1), d_7_-PE33:1(15:0/18:1), d_9_-PE36:1p(18:0p/18:1), d_31_-PS(d_31_-16:0/18:1), d_7_-PA33:1(15:0/18:1), d_7_-PG33:1(15:0/18:1), d_7_-PI33:1(15:0/18:1), C17-SL, _d5-_CL72:8(18:2)4, d_7_-LPE18:1, C17-LPI, C17-LPA, C17-LPS, C17-LPG were obtained from Avanti Polar Lipids. Free fatty acids were quantitated using d_31_-16:0 (Sigma-Aldrich) and d_8_-20:4 (Cayman Chemicals).

### Purification of MVB and ER membranes by density gradient centrifugation

OptiPrep flotation assays were performed to enrich lysosomal membrane fractions according to a standard protocol^71^. Briefly, HeLa cells from four confluent 10-cm dishes were collected and resuspended in 2 ml ice-cold homogenization buffer (250 mM sucrose, 20 mM HEPES-KOH, pH 7.4, 1 mM EDTA, 1 mM phenylmethanesulfonyl fluoride, and complete EDTA-free protease inhibitor), followed by the lysis of the cells in a 7-ml Dounce homogenizer with 15–25 strokes. The homogenized cells were centrifuged twice at 3,000 × g for 10 min to remove cell debris and undisrupted cells. The supernatant was diluted with an equal volume of OptiPrep (D1556; Sigma-Aldrich). A discontinuous OptiPrep gradient was generated in an SW41 tube for ultracentrifuge rotors (344059; Beckman Coulter) by overlaying the following OptiPrep solutions all in homoge-nization buffer: 2.4 ml of the diluted supernatant in 30% Opti-Prep, 1.8 ml 20% OptiPrep, 2 ml 15% OptiPrep, 2 ml 10% OptiPrep, 2 ml 5% OptiPrep, and 2 ml 0% OptiPrep. The gradient was centrifuged at 150,200 × g in an SW41Ti rotor (Beckman Coulter) using an Optima XE-90 ultracentrifuge (Beckman Coulter) for 3 h, and subsequently, 14 fractions (0.85 ml each) were collected from the top and analyzed by Western blots.

ER fractions were enriched using Endoplasmic Reticulum Isolation Kit (ER0100; Sigma-Aldrich) according to the manufacturer’s instructions. Briefly, HeLa cells from five confluent 10-cm dishes were collected, followed by centrifugation at 600 × g for 5 min. After washing the cells three times with PBS, the packed cell volume (PCV) was measured and then suspended in a volume of hypotonic extraction buffer (10 mM HEPES, pH 7.8, with 1 mM EGTA and 25 mM potassium chloride) equivalent to three times the PCV. After the incubation of the cells for 20 min at 4°C allowing the cells to swell, the cells were centrifuged at 600 × g for 5 min, followed by the measurement of the “new” PCV. After adding a volume of isotonic extraction buffer (10 mM HEPES, pH 7.8, with 0.25 M sucrose, 1 mM EGTA, and 25 mM potassium chloride) equivalent to two times the “new” PCV, the suspension was then transferred to a 7-ml Dounce homogenizer, followed by the lysis of the cells with 10 strokes and then the centrifugation of the homogenate at 1,000 × g for 10 min at 4°C. After the transfer of the supernatant to another centrifuge tube, the supernatant was centrifuged at 12,000 × g for 15 min at 4°C, followed by another centrifugation for 60 min at 100,000 × g at 4°C. The pellet was the microsomal fraction and further verified by Western blots using anti-Calnexin antibody.

### Correlative light electron microscopy

Cells were grown on glass-bottom P35G-2-14-C-Grid dishes (MatTek). The dishes have a high optical quality coverslip with a photo-etched grid and coordinates to facilitate pinpointing the location of individual cells. The cells were fixed with 2% paraformaldehyde (PFA, 16% paraformaldehyde, Ted Pella Co.) in 0.1 M PB buffer pH 7.3 for 30 min at RT. Once the cells of interest were found, their positions on the grid were documented by switching from fluorescence to differential interference contrast (DIC) mode. After fluorescence imaging, the selected areas with positive cells were marked on the bottom of the coverslip under a light microscope to facilitate the processing of EM. After observation with a confocal microscope (Zeiss LSM 980), the samples were fixed in 2.5% glutaraldehyde (25% glutaraldehyde ampules, Ted Pella Co.) and 2% PFA mixture in 0.1 M PB buffer pH 7.3 at 4°C overnight. After washing with PB buffer three times, the samples were stained with 2% OsO4 and 1.5% potassium hexacyanoferrate, followed by sequential washing with PB buffer (three times) and ddH2O (three times). Then, the samples were incubated in 1% TCh and washed with ddH2O 4 times, followed by staining with 2% OsO4 along with another washing step with ddH2O four times. The samples were then stained with 1% UA, followed by washing with ddH2O four times. The samples were stained with a lead aspartate solution and washed with ddH2O five times. Then the samples were dehydrated by incubating with ethanol (30, 50, 70, 95, and 100% X2), followed by incubation with acetone two times. After hydration, the samples were subjected to infiltration and embedding step, in which samples were infiltrated with Epon resin (EMS Corp.) as follows: 3:1, 1:1, 1:3 Acetone: Epon and 100% Epon three times, followed by samples being embedded and polymerized with Epon for 48 h at 60°C. Eventually, the samples were applied to the serial ultrathin sectioning step, and the sections were observed at 80 kV in an FEI Talos 120 kv transmission electron microscopy.

### Animal study

To establish a cell line-derived tumor xenograft model, 6-week-old female BALB/c BALB/c mice were randomly assigned to different treatment groups. 4T1 cells (5×10^5 cells suspended in 100 μL PBS) were subcutaneously injected into the right flank of mice. To maximize the supplementation of exosomes in the tumor microenvironment of the BLTP2-depleted group, exosomes derived from wild-type 4T1 cells were subcutaneously injected near the tumor injection site starting from the day after tumor implantation, three times per week, with a total dose of 20 μg. Once the tumor volume reached an average of 50–80 mm³, the injections were switched to intratumoral administration of exosomes. The mice were weighed every two days, and tumor volume was measured using a digital caliper. Tumor volume was calculated using the formula: (length × width²)/2. Mice were euthanized before tumor volume exceeded 1500 mm³.

### Hematoxylin & eosin staining

Tumor tissues harvested from mice were fixed in 4% paraformaldehyde, paraffin-embedded, and sectioned at 5-μm thickness. Sections were deparaffinized, rehydrated, and stained with hematoxylin (5 min) to visualize nuclei, followed by eosin (3 min) to counterstain cytoplasmic and extracellular components. After dehydration and clearing with xylene, slides were mounted with resinous medium.

Histopathological evaluation was performed under a light microscope to assess tumor morphology, necrosis, and stromal architecture.

### Statistical analysis

All statistical analyses and p-value determinations were performed in GraphPad Prism6. All the error bars represent Mean ± SD. To determine p-values, ordinary one-way ANOVA with Tukey’s multiple comparisons test were performed among multiple groups and a two-tailed unpaired student t-test was performed between two groups.

## Acknowledgements

We thank Dr. Kristen Sadler from Scribendi (www.scribendi.com) for editing a draft of this manuscript. We thank the Mass Spectrometry Core Facility (Mr. Cookson K. C. Chiu) and the Bio-imaging Core Facility (Dr. Zhenglong Sun and Ms. Mei Yu) of Shenzhen Bay Laboratory for providing technical supports.

## Author contributions

J. Wang and W. Ji conceived the project and designed the experiments. J. Wang, D. Li, Y. Liu, T. Zhou performed the experiments. J. Wang, D. Li, Y. Liu, J. Xiong and W. Ji analyzed and interpreted the data. W. Ji prepared the manuscript with inputs and approval from all authors.

## Data and materials availability

All the data and relevant materials, including reagents and primers, that supports the findings of this study are available from the corresponding author upon reasonable request.

---

**Fig. S1.**
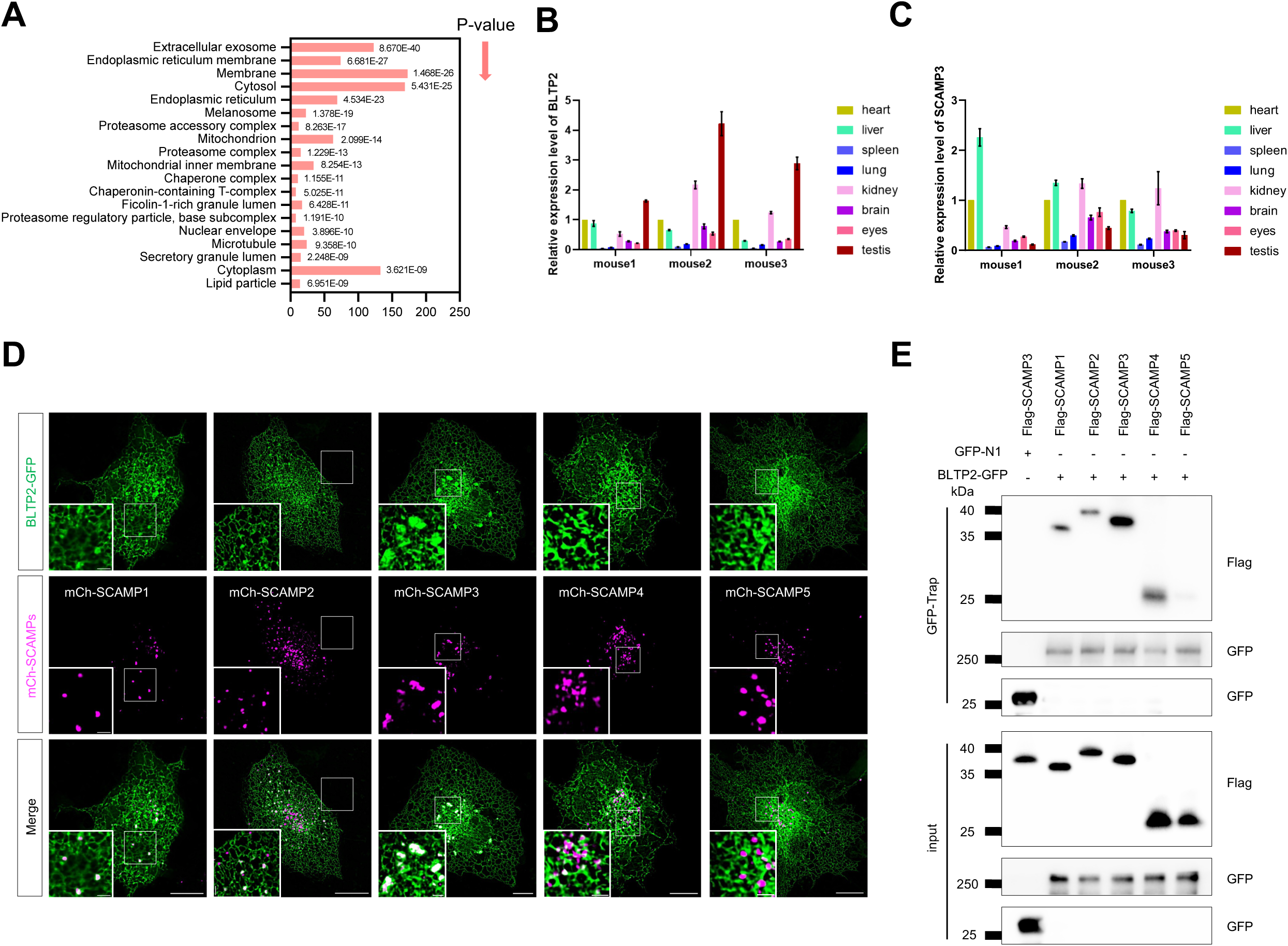
Proteomics analyzes of BLTP2-interacting proteins and the interactions between BLTP2 and other SCAMP proteins. **A:** Proteomics analyzes of BLTP2-GFP interacting proteins in HEK293 cells. **B-C:** The mRNA expression levels of BLTP2(**B**) or SCAMP3(**C**) in different tissues of three mice by qPCR from 3 independent experiments. Mean ± SD. **D:** Representative images of live COS7 cells expressing BLTP2-GFP (green) and mCh-SCAMPs (magenta) with insets. **E:** GFP-Trap assays demonstrate interactions between BLTP2-GFP and mCh-SCAMPs in HEK293 cells. Scale bars: 10 μm in the whole cell images and 2 μm in the insets in (D).

**Fig. S2.**
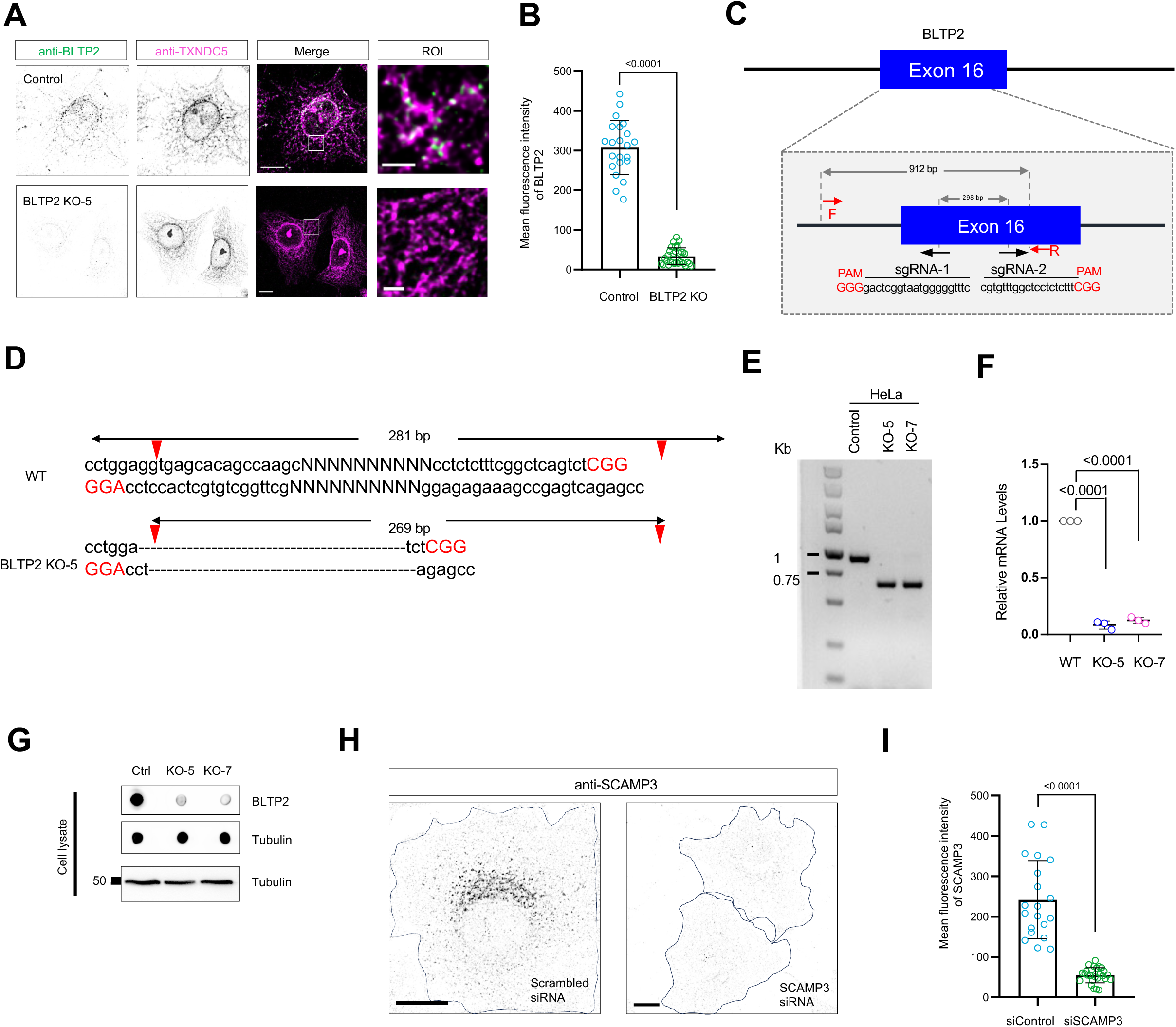
BLTP2 KO HeLa cells by CRISPR-Cas9 and validation of BLTP2 and SCAMP3 antibodies in IF. **A:** Representative images of fixed control or BLTP2 KO-5 HeLa cells stained with antibodies against BLTP2 (green) and TXNDC5 (magenta) with insets on the right. **B:** Mean fluorescence intensity of anti-BLTP2 in control or BLTP2 KO-5 as in (**A**) from three independent experiments. Two-tailed unpaired Student’s t test. Mean ± SD. **C:** CRISPR knock-out of BLTP2 in HeLa cells (BLTP2-KO). **D:** Genomic sequencing of BLTP2 KO-5 HeLa cells. **E-G:** Two BLTP2 KO clones (#5 and #7) were confirmed by DNA gels (**E**), qPCR (**F**) and dot blot (**G**). **H:** Representative images of fixed HeLa cells stained with antibodies against SCAMP3 upon scrambled or SCAMP3 siRNA. **I:** Mean fluorescence intensity of endogenous SCAMP3 in scrambled or SCAMP3 siRNA cells in more than three independent experiments as in (**H**). Two-tailed unpaired Student’s t test. Mean ± SD. Scale bars: 10 μm in the whole cell images and 2 μm in the insets in (A & H).

**Fig. S3.**
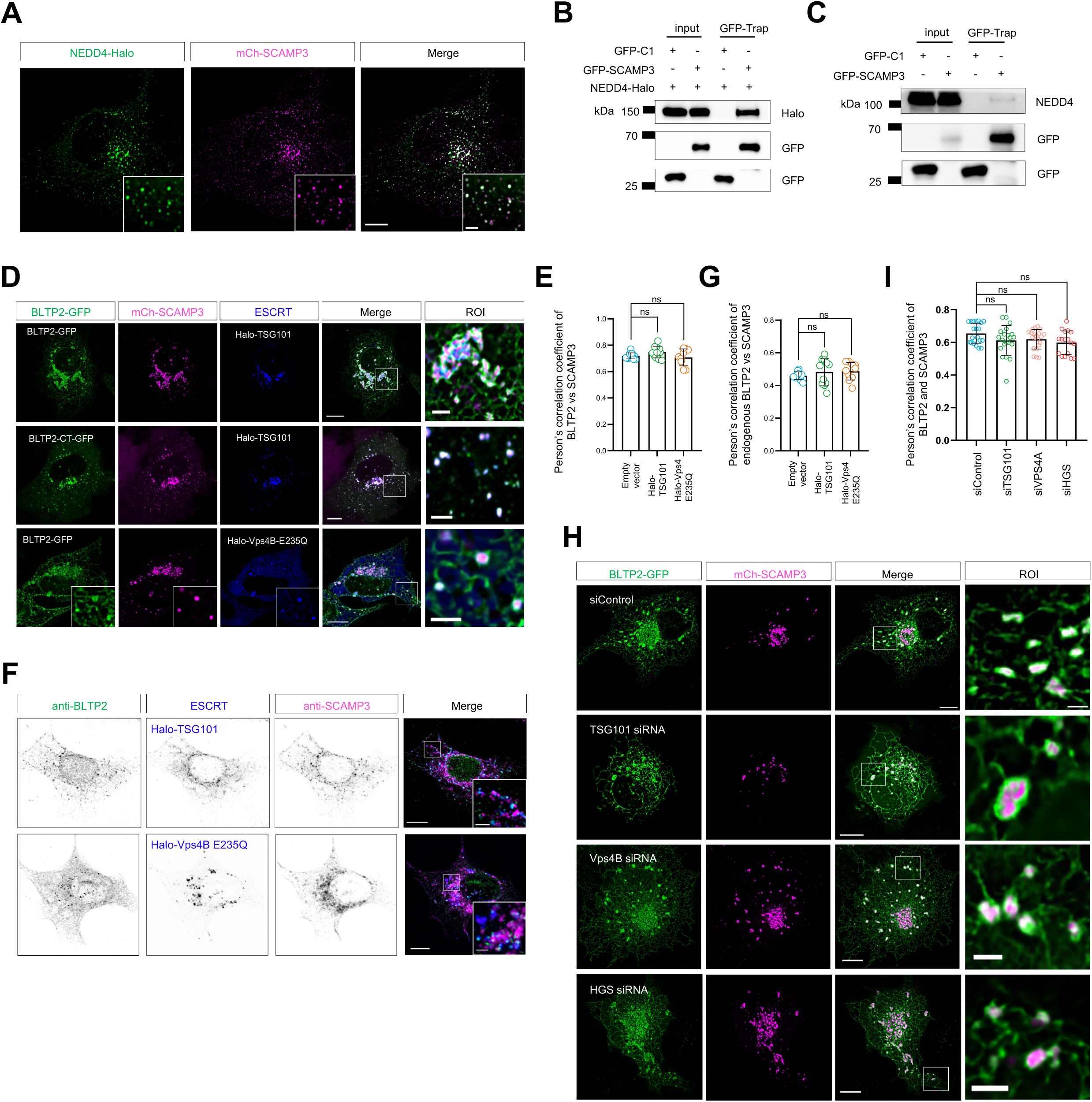
BLTP2 and SCAMP3 associate with the ESCRT complex. **A:** Representative images of a COS7 cell expressing NEDD4-Halo (green) and mch-SCAMP3 (magenta) with insets. **B:** Co-IP assays show interactions between GFP-SCAMP3 and NEDD4-Halo in HEK293 cells. **C:** Co-IP assays show interactions between GFP-SCAMP3 and endogenous NEDD4 in HEK293 cells. **D:** Representative images of live COS7 cells expressing either BLTP2-GFP or BLTP2-CT-GFP (green), mCh-SCAMP3 (magenta) and ESCRT proteins (TSG101 or Vps4B-E235Q; blue) with insets on the right. **E:** Pearson’s correlation coefficient of BLTP2-GFP vs mCh-SCAMP3 in the presence of empty vector, Halo-TSG101 or Halo-Vps4(E235Q) in more than three independent experiments as in (**D**). Ordinary one-way ANOVA with Tukey’s multiple comparisons test. Mean ± SD. **F:** Representative images of fixed HeLa cells stained with BLTP2 antibody (green) and SCAMP3 antibody (magenta) with insets. **G:** Pearson’s correlation coefficient of BLTP2 vs SCAMP3 in the presence of empty vector, Halo-TSG101 or Halo-Vps4(E235Q) in more than three independent experiments as in (**F**). Ordinary one-way ANOVA with Tukey’s multiple comparisons test. Mean ± SD. **H:** Representative images of live COS7 cells expressing BLTP2-GFP (green) and mCh-SCAMP3 (magenta) treated with either TSG101, VPS4B or HGS siRNAs with insets on the right. **I:** Pearson’s correlation coefficient of BLTP2 vs SCAMP3 in scrambled, TSG101, VPS4A or HGS siRNA-treated cells in more than three independent experiments as in (**H**). Ordinary one-way ANOVA with Tukey’s multiple comparisons test. Mean ± SD. Scale bars: 10 μm in the whole cell images and 2 μm in the insets (A, D, F & H).

**Fig. S4.**
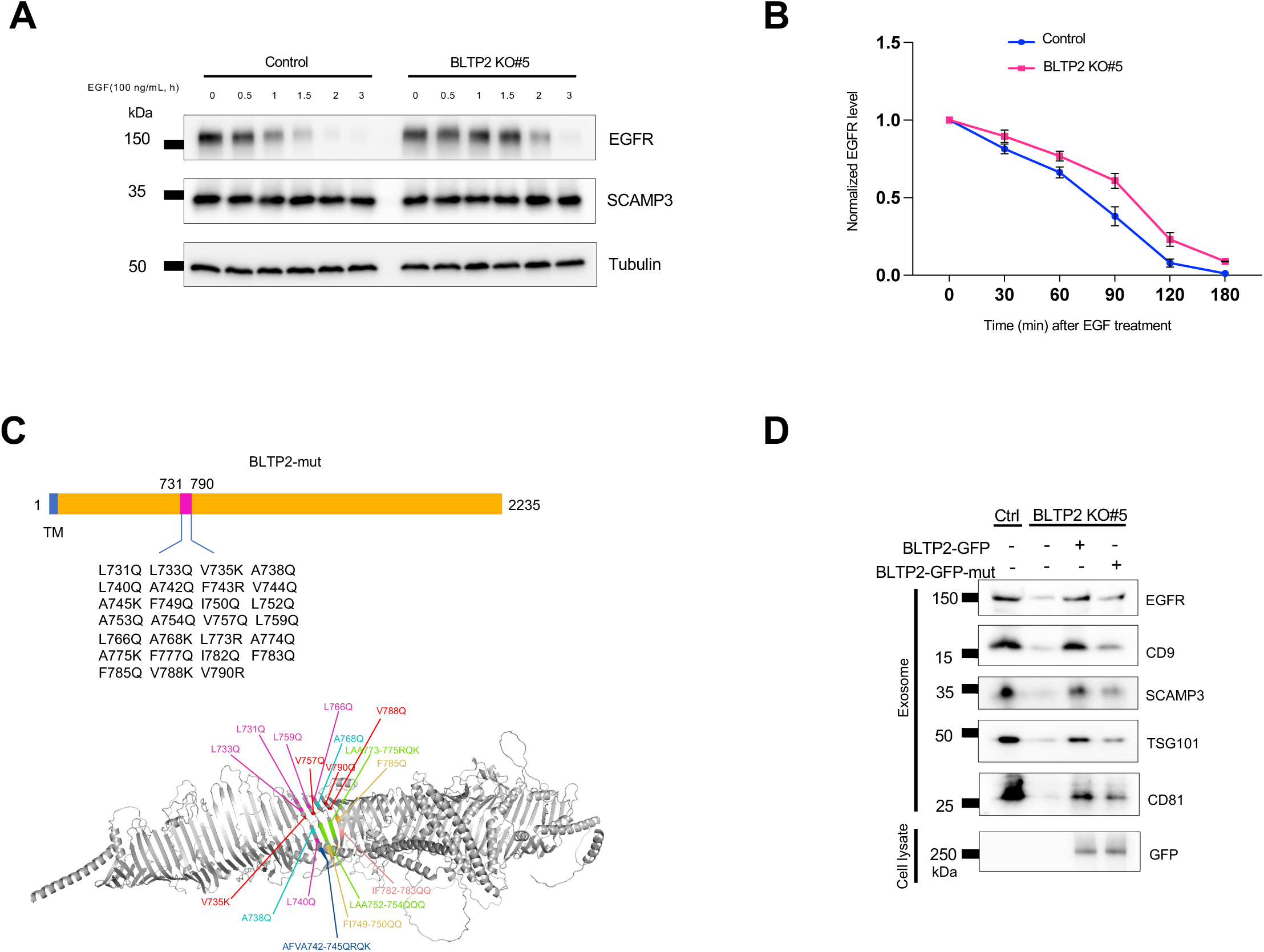
BLTP2 promotes exosome formation depending on its lipid transfer activity. **A:** Western blotting showing the EGFR levels in control or BLTP2 KO-5 HeLa cells at different time points after EGF (100 ng/mL) treatment. **B:** Quantitation of EGFR levels in three independent experiments as in (**A**). The EGFR levels were normalized to time 0. Mean ± SD. **C:** Schematic diagram of the lipid transfer-deficient BLTP2 mutant. **D:** Representative Immunoblotting of the purified exosomes from the control, BLTP2 KO, or BLTP2 KO expressing either WT BLTP2-GFP or the lipid transfer-deficient mutant.

**Fig. S5.**
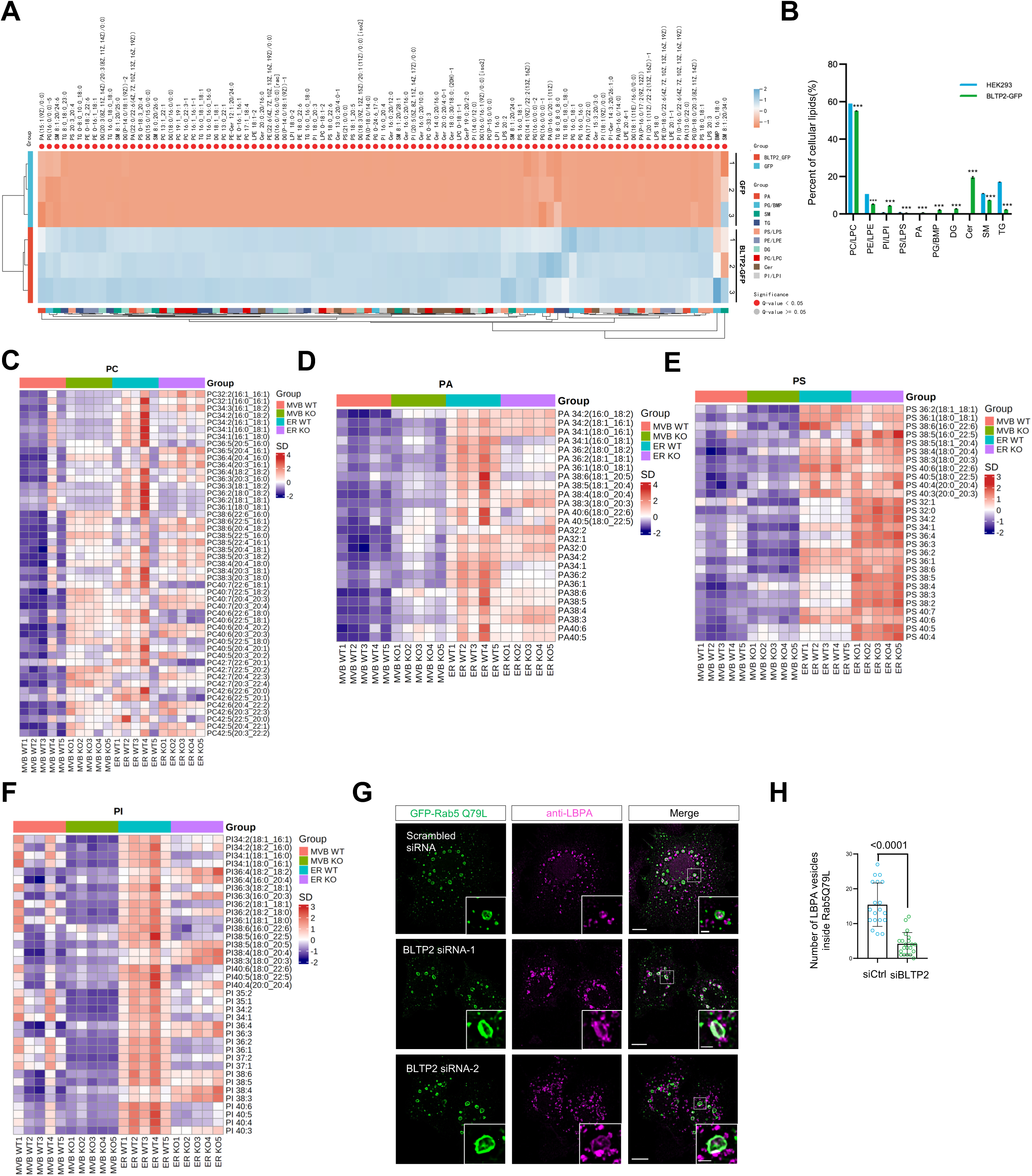
BLTP2 associates with glycerophospholipids such as PE and PG. **A:** Heatmap showing different lipid species associated with GFP vector or BLTP2-GFP in HEK293 cells in three independent experiments. **B:** Percentage of lipids that were associated with BLTP2–GFP (green) and cellular abundance of these glycerolipids^50^ is indicated in blue. **C-F:** The levels of different lipid species of PC (**C**), PA (**D**), PS (**E**) and PI (**F**) in the endosomal fraction or the ER fraction from control or BLTP2 KO-5 HeLa cells in five independent experiments as in Fig. 7A**-C**. **G:** Representative images of fixed HeLa cells expressing GFP-Rab5-Q79L (green) and stained with LBPA antibody (magenta) upon scrambled or two BLTP2 siRNAs with insets. **H:** The number of LBPA vesicles inside Rab5-Q79L endosomes upon scrambled or BLTP2 siRNAs in more than three independent experiments as in (**G**). Two-tailed unpaired Student’s t test. Mean ± SD. Scale bars: 10 μm in the whole cell images and 2 μm in the insets (G).

